# Effects of hypoxia and low temperature on female physiology and reproduction of *Drosophila melanogaster*

**DOI:** 10.64898/2026.04.08.717251

**Authors:** N Rivera-Rincón, EC Saurette, AE May, AG Appel, LS Stevison

## Abstract

Because hypoxia and low temperature independently alter metabolism and reproductive investment, their interaction provides a tractable framework for testing whether combined stressors produce non-additive physiological and reproductive effects. Here, we investigated the single and combined effects of hypoxia and low temperature in *Drosophila melanogaster* across multiple genetic backgrounds. We quantified metabolic rate, thermal tolerance, body mass, fertility, oogenesis progression, and oocyte apoptosis to assess organismal responses to environmental stress. Hypoxia generally increased respiratory quotient and body mass, but its effects on thermal tolerance and fertility were highly genotype dependent. Across traits, combined stressors frequently produced responses that differed from those observed under single stressors, including reduced fertility, altered oogenesis, and changes in oocyte cell death. Importantly, these effects were not uniform: some genotypes exhibited increased oocyte production or reduced cell death under combined stress, highlighting pronounced genotype-dependent differences in stress sensitivity and reproductive allocation. Together, our results demonstrate that the interaction between hypoxia and temperature can modulate metabolic and reproductive responses in ways that are not predictable from single-stressor responses alone. These findings highlight the importance of incorporating genetic background and interacting environmental stressors when evaluating organismal tolerance and adaptive potential under ongoing environmental change.

## 1. Introduction

Understanding how multiple stressors affect living organisms is critical to predicting their resilience and plastic responses to changing environments (Gunderson et al., 2017; Helmuth et al., 2014). Most studies focus on single stressors, yet organisms in nature are typically exposed to multiple, interacting stressors that can enhance, mitigate, or neutralize one another’s effects (Crain et al., 2008; Gunderson & Stillman, 2015; Piggott et al., 2012). These interactions challenge plasticity capacities by combining physiological and ecological pressures. For example, shifts in temperature and oxygen availability often occur simultaneously, demanding a better understanding of their combined impacts on behavior, physiology, and survival (Boyd et al., 2018; Folguera et al., 2011).

Reproductive functions are particularly vulnerable to environmental stress, as energy-intensive processes such as oogenesis and egg-laying are directly affected by metabolic and physiological changes (Folguera et al., 2011; Rivera-Rincón et al., 2024; Zizzari & Ellers, 2014). In females, these stress-induced demands can compromise reproductive output and fitness, influencing population dynamics (Lord et al., 2021; Meiselman et al., 2018; Teder & Kaasik, 2023). Given the ecological importance of reproduction, understanding how combined stressors impact female fitness provides insights into broader ecological consequences. Studying the impact of combined stressors is especially biologically relevant for natural populations, where multiple stressors often co-occur and present unique challenges to reproduction (Boyd et al., 2018; Gunderson et al., 2017).

The model organism *Drosophila melanogaster* offers valuable insights into stress responses, as it is well-studied under single stressors, such as hypoxia and extreme temperatures (T. Gorr et al., 2006; T. A. Gorr et al., 2004; Hoffmann et al., 1997). Drosophila larvae are naturally adapted to hypoxic conditions in fermenting fruits and thus, exhibit physiological resilience that highlights the role of genetic background in coping with environmental stress (Lee et al., 2019; Valko et al., 2022). However, the interaction between stressors such as hypoxia and cold remains underexplored, as does the extent to which genetic variability influences these responses. In *Drosophila*, different genotypes exhibit varied responses to individual stressors such as hypoxia or cold, with some showing heightened resilience and others demonstrating significant declines in performance (Lazzaro et al., 2008; Mackay et al., 2012; Singh, 2019; Weber et al., 2012). Certain genotypes may exhibit synergistic effects, where combined stressors amplify physiological or reproductive declines, while others demonstrate tolerance or compensatory responses. Incorporating genetic background into studies of environmental stress can improve predictions of population responses and their long-term resilience to multifaceted ecological challenges (Flatt, 2011; Hoffmann & Sgrò, 2011).

This study aimed to describe the effects of hypoxia and low temperature individually and their interaction on both the physiological and reproductive traits of *Drosophila*. Key metrics, including Respiratory Quotient, Critical Thermal Maximum, and body mass, were assessed alongside reproductive success metrics like fertility and oogenesis progression. By examining genetic variability’s role in responses to combined stressors, this research aims to better understand how genetic variation shapes physiological and reproductive plasticity under interacting environmental stressors. These findings are critical for understanding resilience and plasticity in natural populations, with implications for understanding how genotype-by-environment interactions shape responses to global climate change.

## 2. Methods

### 2.1. Genetic Background

The *Drosophila* Genome Reference Panel (DGRP) (Huang et al., 2014) consists of over 200 fully sequenced lines derived from a natural population in North Carolina (Mackay et al., 2012). These lines exhibit extensive phenotypic and genotypic diversity across a wide range of traits. From this panel, we focused on two previous experimental manipulations related to our perturbations. While no study manipulated oxygen concentration directly, a previous study treated the flies with a drug known as paraquat through their diet, to induce oxidative stress and measured survival across all 200 lines (Weber et al., 2012). These two stressors are known to have overlapped genetic and molecular pathways, with hypoxia generating ROS and subsequent oxidative damage as a consequence of low oxygen (Zhao and Haddad 2012). A second study measured chill coma recovery time following 3 hours under ice (Mackay et al., 2012). Based on the results from these two studies, we selected a subset of five DGRP lines that had a range of paraquat survival and chill coma recovery times representing the distribution of values across each experiment. Specifically, we selected lines DGRP-42 (BDSC_25193), DGRP-57 (BDSC_29652), DGRP-391 (BDSC_25191), DGRP-491 (BDSC_28202), and DGRP-508 (BDSC_28205), which were obtained from the Bloomington *Drosophila* Stock Center with Research Resource Identifiers in parentheses for each stock (see Supplemental figure S1).

### 2.2. Treatments

To select a non-lethal stressful low temperature, we conducted pilot experiments and compared fertility for a different DGRP-217 (BDSC_28154) line at 14℃, 16℃, and 18℃. The temperature that resulted in the highest mean per female across these temperatures (18℃) was selected for subsequent experiments as the low temperature (see Supplemental figure S2).Because this pilot was conducted on DGRP-217, which was not included in the main experiment, the optimal low temperature may differ for the focal lines given the genotype-dependent responses observed throughout this study. Additionally, 25℃ was used as a control temperature since it is the standard temperature used in *Drosophila melanogaster* rearing in laboratory environments. Oxygen availability was selected based on previous studies on survivorship at different oxygen concentrations (Zhou et al., 2007). Specifically, we selected 8% oxygen throughout development, which was where the egg-to-adult survivorship was comparable with the temperature treatments (survival reduction of 20%). We selected oxygen levels of 21% O_2_ and 8% O_2_ as normoxia and hypoxia, respectively. Custom-made chambers (BioSpherix) were used to control oxygen levels and were placed into incubators to control temperature. Certified tanks from AirGas, with either 21% or 8% oxygen, were used to flush the chambers for twenty minutes, every 12 hours, and every time the chambers were opened. Each chamber had a small USB powered fan attached, to help with the air circulation within the chamber.

### 2.3. Fly husbandry

All lines were maintained at ± 25°C with a photoperiod of 12:12 (light:dark) in an incubator (Percival DR41-VL). Flies were reared on standard cornmeal-sugar-yeast-agar media (Bloomington *Drosophila* Stock Center recipe) in polypropylene enclosures. Stocks were routinely authentication via a RFLP-based protocol using custom markers for these stocks (Shiran et al., 2026).

### 2.4. Genetic Crosses

A total of 200 genetic crosses were set up, 20 per genetic background per treatment. An average of 10 mutant females and 5 DGRP males age-matched (2 days old, post eclosion from pupal case) were used for the crosses. These genetic crosses were allocated to one of the four different treatment combinations: control (Normoxia + Control temperature), hypoxia (Hypoxia + Control temperature), low temperature (Normoxia + Low temperature), or combined (Hypoxia + Low temperature). The F_1_ generation was then measured across six parameters: Respiratory Quotient (RQ), Critical Thermal Maximum (CT_max_), body mass, fertility, stages of oogenesis, and cell death (Fig. 1).

**Figure 1:**
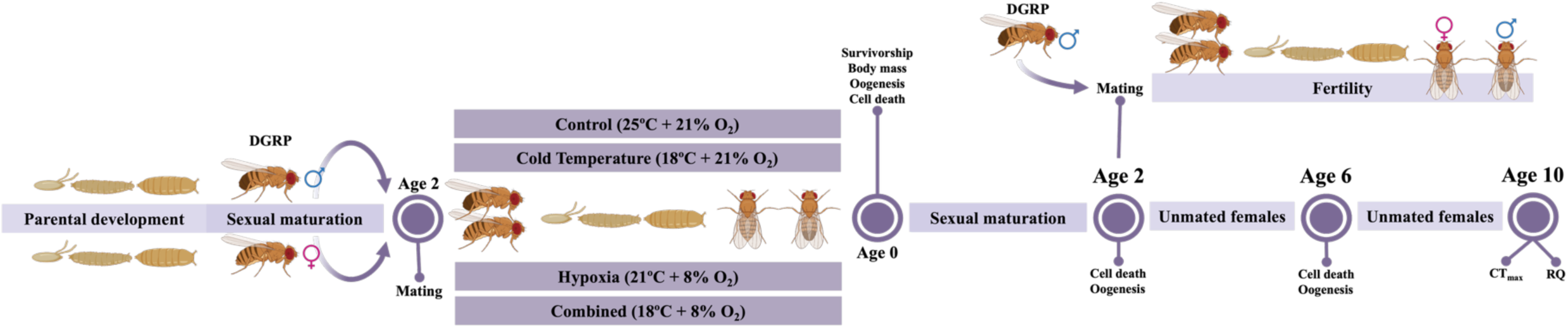
Graphical representation of the experimental design, with the ages at which each measurement was taken. Age is defined as time post-eclosion from pupal case in days.

### 2.5. Physiology

Respiratory Quotient (RQ). The respiratory quotient (RQ) was calculated as the ratio of CO₂ produced to O₂ consumed. In this study, both CO₂ and O₂ were measured in individual flies at 10 days post-eclosion. This age was selected because, as previous research shows, metabolic rates remain relatively stable in *Drosophila melanogaster* at this stage (Arking et al., 1988; Hulbert et al., 2004; Mockett et al., 2001; Promislow & Haselkorn, 2002; Van Voorhies et al., 2003, 2004). RQ was determined using closed-system respirometry following the method outlined by DeVries and Appel (2013). Briefly, flies were weighed both before and after incubation. Each fly was placed in an individual 1 mL syringe serving as a respirometry chamber, flushed with dry, CO₂-free air, sealed, and incubated in the dark at the treatment temperature for 4 hours (and no longer than 5 hours, to avoid starvation stress).

After incubation, a 0.25 mL air sample from each chamber was injected into a respirometry system for analysis, and data was collected with *ExpeData* software (Sable Systems, North Las Vegas, NV, USA). Calculations involved converting the recorded data into units of mL/min, integrating the CO₂ and O₂ peaks, and standardizing by body mass to determine total CO₂ production or O₂ consumption per chamber. RQ was then calculated as the ratio of CO₂ produced to O₂ consumed. To ensure accuracy, metabolic rates were measured for at least 10 female flies per treatment per genetic background. Any RQ values exceeding 1.5, which was associated with flies that appeared dead during measurement, were excluded from analysis (n = 20).

Critical thermal maximum (*CT_max_*). Ten-day-old flies were tested in a custom-designed, microprocessor-controlled incubator that maintained a constant heating or cooling rate (Hu & Appel, 2004). Each fly was individually weighed and placed in a small chamber inside the incubator, along with an additional open chamber containing water to minimize desiccation. The temperature was gradually increased at a rate of 0.1°C per minute, and flies were monitored for knockdown. CT_max_ was defined as the highest temperature at which a fly was knocked down, unable to right itself, but still able to recover afterward, following the methods of Sponsler and Appel (1991). CT_max_ measurements were taken from at least 15–20 females per treatment, and genetic background.

Body mass was measured in freshly eclosed females (age 0 days post-eclosion) to capture whole-organism mass immediately following development under each treatment. Females were anesthetized with CO₂, placed individually in pre-tared weighing boats, and weighed once each. Body mass was recorded to the nearest 0.00001g. The tare function in the balance was used before and after each individual measurement. All weights were taken using a Sartorius 1712 MP8 balance. A minimum of 10 females per treatment and genetic background were used for these measurements. This metric was included as an additional physiological measure to assess treatment associated differences in whole-organism mass allocation following development under hypoxia and temperature treatments.

### 2.6. Reproduction

F_1_ fertility. For 7 days, all flies were cleared from each cross vial, and the number of adult females and males was recorded. F_1_ fertility was calculated by dividing the total adult offspring per vial by the number of females used in the cross.

F_2_ Fertility. Ten virgin females from the F_1_ crosses were mated with 5 males from the same DGRP line used on the F_1_ cross (Fig. 1). These F_2_ crosses were placed into an incubator and reared at 25°C. The crosses were transferred into new vials every 2 days for 8 days (A, B, C, D), and the number of adults was recorded daily for 7 days for each vial. F_2_ fertility was calculated by dividing the total adult offspring across the 8 days by the number of females used in the cross.

Oogenesis Progression and Apoptosis. Using previous studies by Rivera-Rincón et al. (2024), three time points (age in days post-eclosion of pupal case) were selected to compare the differences in oogenesis and apoptosis due to changes in temperature and oxygen availability: early (age 0), mid (age 2), and late (age 5). The early time point consisted of only oocytes at the germarium stage. Mid-time point was defined as consisting of oocytes in all stages, with its majority at developmental stages 1-7. The late time point was defined as consisting of oocytes in all stages with its majority at the developmental stages 12-14 (Fig. 2). Dissections of the adult flies were performed in the morning for consistency across time points and samples, ensuring that no more than 20 minutes passed between the first dissection and its fixation to avoid tissue degradation. Following the protocol Meehan et al. (2015) described, the tissue was stained with the fluorescent stain DAPI (Vectashield with DAPI) for nuclei labeling. Fluorescein-12-dUTP was used via the DeadEnd™ Fluorometric TUNEL System from Promega to detect DNA degradation. High-resolution images were captured using a BZ-X fluorescence microscope.

**Figure 2:**
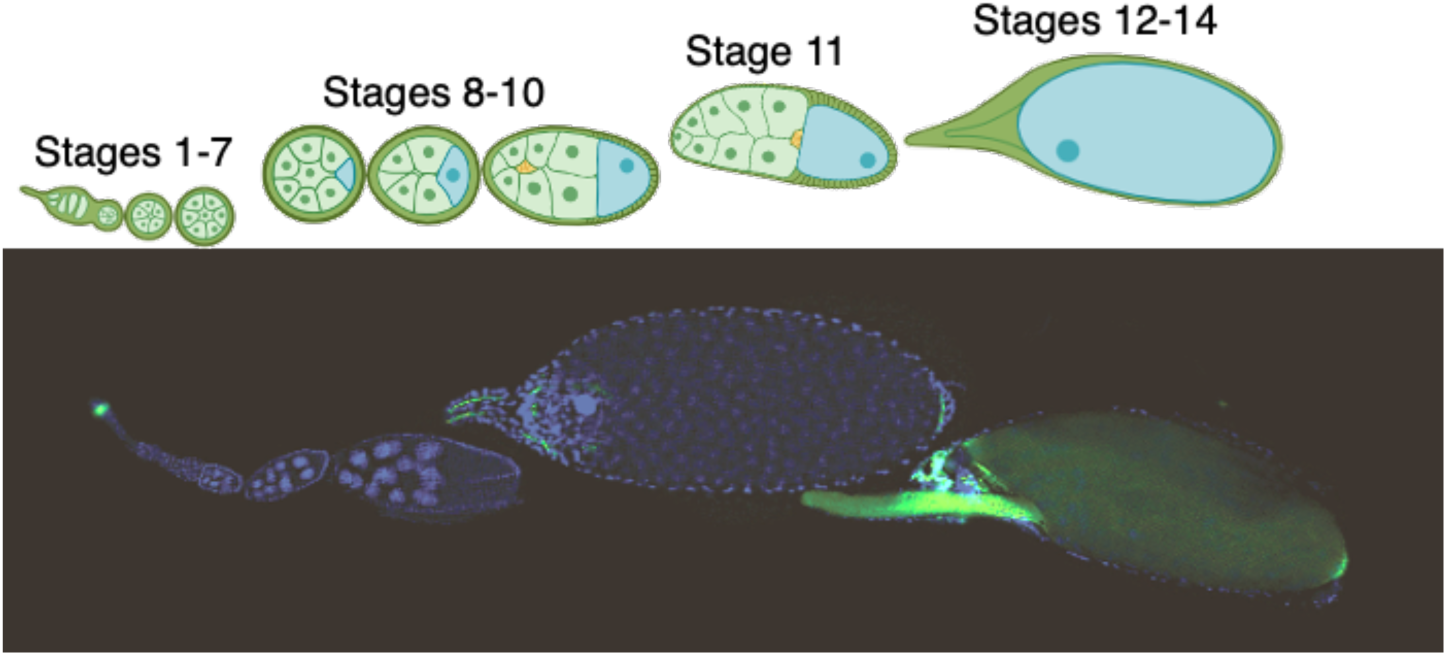
Redrawn of the egg chamber’s developmental stages from Rivera-Rincón et al.,2024, with composite image from this study.

### 2.7. Image Analysis

Using the oocyte developmental stages described by Jia et al. (2016), oocytes were pooled into five developmental time points: Germarium, stages 1 – 7, stages 8 – 10, stage 11, and stages 12-14 (Fig. 2). Through the Jupyter platform (Kluyver et al., 2016), the Scikit-image (v0.22.0) (Van der Walt et al., 2014) and the Napari (Chiu et al., 2022) packages were used to standardize each DAPI image in terms of size and brightness and transform it to black and white for better nuclei identification. Annotation of the images was performed using a trained AI model through the APEER platform (Dang et al., 2021), which was then used to identify, characterize, and count oocytes from the image data. At least 50 objects from each class (developmental stages and background) were individually annotated in a randomly selected subset of images for AI model training. The model was updated with new and corrected labels on each iteration until the detection accuracy exceeded 95%. The AI model was then used to create an instance segmentation for each image, to detect and identify individual objects (characterize), and to segment them as separate instances (quantify). The segmentation was inspected, and images with poor characterization due to blurry backgrounds or overlapping objects that impeded accurate characterization were removed (n=3). The number of oocytes per stage was standardized with the number of ovarioles present per ovary. Using the extension “Fiji” of the software, ImageJ (Schindelin et al., 2012), each image was split into three channels, and using the third (green) channel only, TUNEL brightness was used to determine the presence or absence of DNA-fragmentation per oocyte.

### 2.8. Statistical analysis

All statistical analyses were conducted in R v4.0.1. Genetic background was treated as a fixed effect, aiming to focus on the contrasting response profiles among a defined subset of DGRP lines selected *a priori* for divergent stress responses. Physiological measurements and F_1_ fertility were analyzed using a linear mixed-effects models fitted with the ‘lmer’ function from the “lme4” package (v.1.1-34, Bates et al. (2014)). Oxygen availability, temperature, and genetic background were included as fixed effects along with their interactions. Vial was included as a random effect to account for non-independence among individuals reared together (*e.g., Physiology (1|vial) + Oxygen*Temperature* Genetic Background*). For F_2_ fertility, mixed-effects models were also used, with vial number included as a random effect and day as a blocking factor to account for temporal variation in offspring counts (*e.g., Fertility (1|vial) + (1|Day) + Oxygen*Temperature* Genetic Background*).

The number of oocytes was analyzed using a zero-inflated negative binomial mixed-effects model, which is appropriate for overdispersed count data with excess zeros. The model was implemented using the R package ‘glmmTMB’ v1.1.9 (Brooks et al. 2017; McGillycuddy et al. 2025). The model included fixed effects for Stage, Timepoint, Temperature, Oxygen, and Stock, as well as their relevant interaction terms. Ovary identity was included as a random intercept, to account for non-independence among observations from the same ovary, and the zero-inflation component was modeled as a function of Stage to allow the probability of structural zeros to vary across developmental stages. Oocyte cell death (TUNEL assay) was analyzed using a binomial generalized linear mixed-effects model with a logit link function. The response was modeled as the number of TUNEL-positive and TUNEL-negative oocytes. Fixed effects included stage group (early vs late oogenesis), temperature, oxygen availability, genetic background, and their interactions, with ovary identity included as a random intercept. For all models, significance of fixed effects was assessed using Type II Wald χ² tests implemented in the *car* package (Fox et al., 2012). Post hoc comparisons were conducted using estimated marginal means from the *emmeans* package (v. 1.5.5-1, Lenth and Lenth (2018)), with contrasts constructed within specific environmental and genetic contexts and p-values adjusted for multiple comparisons where appropriate. All the figures were generated using “ggplot2” (Wickham, 2011), and “ggpubr” (v.0.6.6, Kassambara and Kassambara (2020)). Final figures schematics were created using Biorender.

### 2.9. Data Availability

All raw data and code are included as part of a public GitHub repository: https://github.com/StevisonLab/Combination-effects-of-hypoxia-and-low-temperature-in-Drosophila/. We plan to make a static version upon acceptance and include a DOI here. Also, the ovary images will be deposited in the Aurora Institutional Repository upon acceptance.

## 3. Results

A total of 3,487 females were used to measure F_1_ fertility. From the 29,668 flies collected across 366 replicates and treatments combinations, 1,680 females were used to measure respiratory quotients (RQ), critical thermal maximum (*CT_max_*), body mass, and F_2_ fertility. In addition, 453 individual ovaries were dissected across treatments, time points, and genetic backgrounds (GB’s) to measure oogenesis and oocyte cell death (see Supplemental Table S1). For clarity, we refer to the four experimental conditions as control, hypoxia only, low temperature only, and combined treatment. Because oxygen availability and temperature were analyzed as crossed factors, responses under the combined treatment correspond to interaction effects between these factors.

### 3.1. Physiology

Respiratory Quotient (RQ). A total of 183 individuals were used to measure metabolic rates across treatments and genetic backgrounds (see Supplemental Table S1). DGRP-391 was excluded from these analyses due to insufficient individuals in the control treatment group, and 20 additional samples were excluded as outliers because they appeared dead during measurement. CO_2_ production, O_2_ consumption, and the respiratory quotient were analyzed separately (Table 1).

**Table 1:**
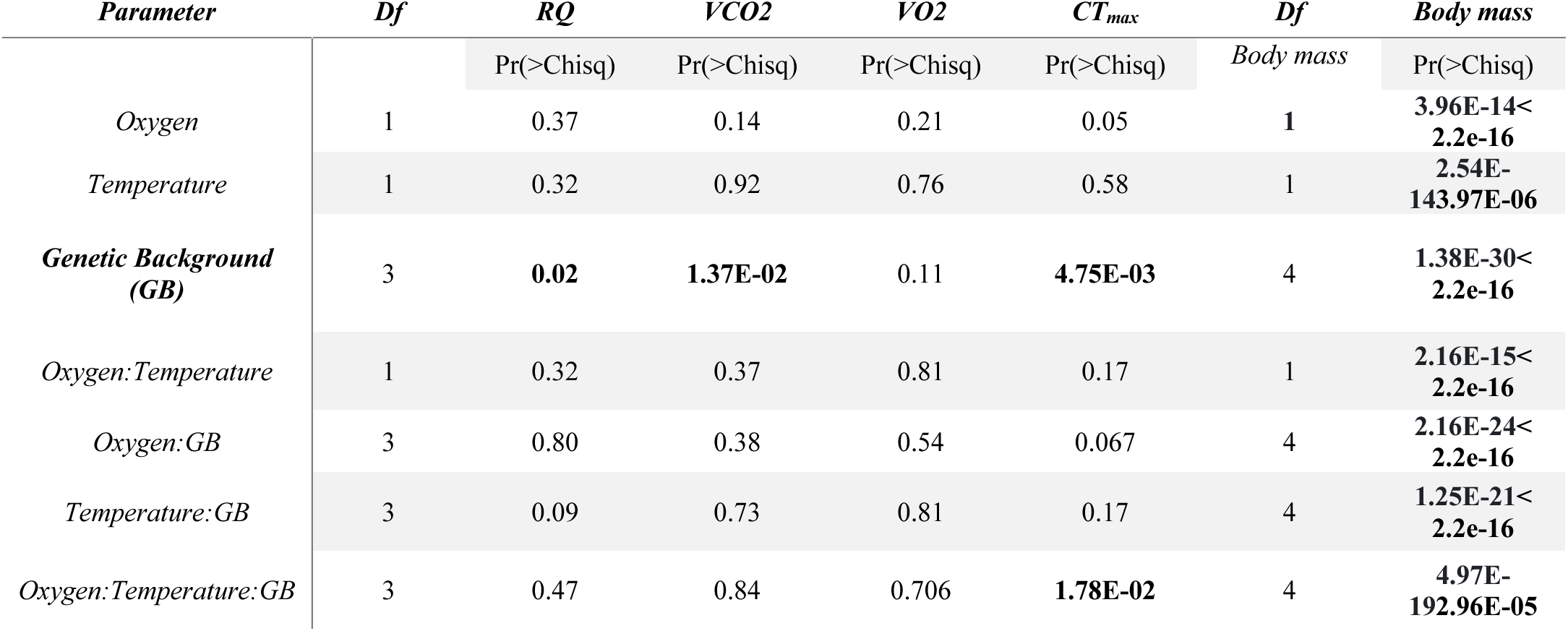
Mixed-effects model results for the effects of oxygen, temperature, genetic background, and their interactions on all the physiological traits measured.

| *Parameter* | *Df* | *RQ* | *VCO2* | *VO2* | *CT<sub>max</sub>* | *Df* | *Body mass* |
| --- | --- | --- | --- | --- | --- | --- | --- |

|  |  | Pr(>Chisq) | Pr(>Chisq) | Pr(>Chisq) | Pr(>Chisq) | Body mass | Pr(>Chisq) |
| --- | --- | --- | --- | --- | --- | --- | --- |
| Oxygen | 1 | 0.37 | 0.14 | 0.21 | 0.05 | 1 | 3.96E-14<2.2e-16 |
| Temperature | 1 | 0.32 | 0.92 | 0.76 | 0.58 | 1 | 2.54E-143.97E-06 |
| Genetic Background (GB) | 3 | 0.02 | 1.37E-02 | 0.11 | 4.75E-03 | 4 | 1.38E-30<2.2e-16 |
| Oxygen:Temperature | 1 | 0.32 | 0.37 | 0.81 | 0.17 | 1 | 2.16E-15<2.2e-16 |
| Oxygen:GB | 3 | 0.80 | 0.38 | 0.54 | 0.067 | 4 | 2.16E-24<2.2e-16 |
| Temperature:GB | 3 | 0.09 | 0.73 | 0.81 | 0.17 | 4 | 1.25E-21<2.2e-16 |
| Oxygen:Temperature:GB | 3 | 0.47 | 0.84 | 0.706 | 1.78E-02 | 4 | 4.97E-192.96E-05 |

RQ differed significantly among genetic backgrounds, but neither oxygen availability nor temperature had a significant overall effect (Table 1). Despite this, the direction and magnitude of treatment responses varied among lines. Under hypoxia, several genetic backgrounds showed modest increases in RQ relative to normoxia, with differences of up to about 0.12. In contrast, DGRP-57 and DGRP-491 showed only slight increases, ∼ 0.01 under low temperature and ∼0.03 under hypoxia, respectively.

A similar genotype-specific pattern was observed for CO₂ production. VCO₂ differed among genetic backgrounds but did not show a consistent overall response to treatment (Table 1). In several lines, hypoxia reduced VCO₂ relative to normoxia by as much as ∼0.55 µl CO₂ g⁻¹ h⁻¹, whereas DGRP-508 showed an increase of up to ∼0.12 µl CO₂ g⁻¹ h⁻¹ under hypoxia. Estimated marginal means values and standard deviations for VCO₂ and VO₂ across treatments and genetic backgrounds are provided in Supplemental Table S2-3. Thermal limits (*CT_max_*). A total of 186 individuals were used to measure *CT_max_* (Table 1). As in the metabolic analyses, DGRP-391 was because of insufficient individuals in the control group (n=35). CT_max_ differed among genetic backgrounds, and treatment effects depended on the interaction between genetic background, oxygen availability, and temperature (Table 1), indicating that thermal tolerance responses were highly line specific. Across most genetic backgrounds, hypoxia was associated with slightly higher CT_max_ values than normoxia, with increases of up to ∼0.4°C. DGRP-491 was a notable exception, showing a pronounced decrease of ∼0.7°C under hypoxia. Responses at low temperature were more variable: DGRP-491 and DGRP-508 had slight increases in *CT_max_*, approximately 0.04°C and 0.16°C, respectively, whereas DGRP-57 and DGRP-42 had decreases of approximately 0.13°C and 0.71°C, respectively (Fig. 3). Overall, treatment-associated shifts in *CT_max_* spanned nearly 1°C across genetic backgrounds. Estimated marginal means values and standard deviations for *CT_max_* across treatments and genetic backgrounds are provided in Supplemental Table S4. Body mass. A total of 196 individuals were independently weighed at eclosion to measure body mass (see Supplemental S1). Body mass differed significantly among genetic backgrounds and was also affected by oxygen availability and temperature, with a significant three-way interaction indicating that responses to treatment differed among genotype (Table 1).

**Figure 3:**
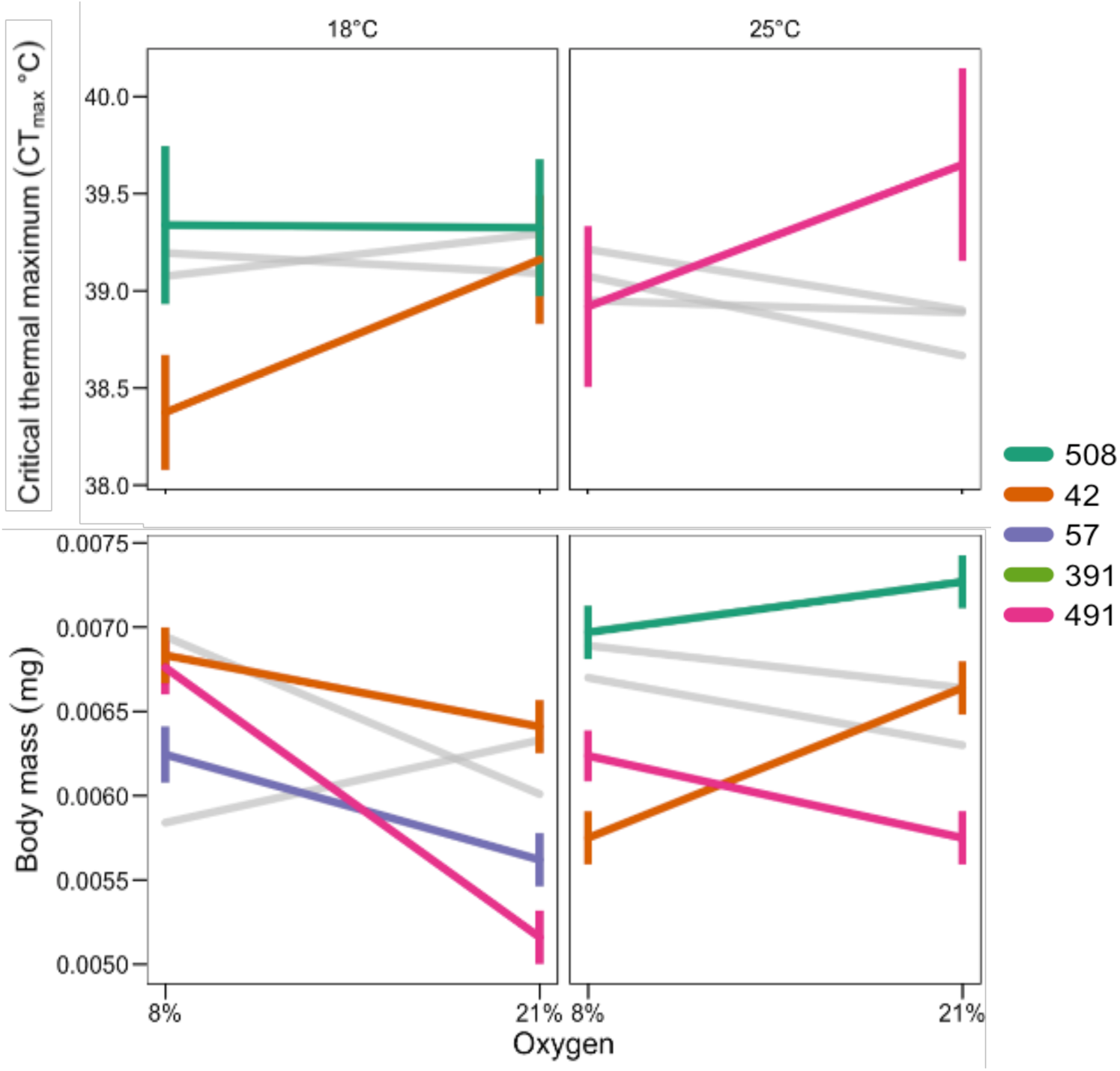
Reaction norm showing the estimates of CT_max_ and body mass measurements for both temperature treatments across two oxygen levels. Stocks are color-coded, with non-gray lines indicating statistical significance from the post-hoc analysis.

Under low temperature, hypoxia was associated with increased body mass in most genetic backgrounds, with increases of up to ∼0.9 mg relative to normoxia. DGRP-391 differed from this general pattern, showing a decrease of approximately 0.4 mg under the same conditions. Under hypoxia alone, most lines showed modest increases in mass (up to ∼0.4 mg), whereas DGRP-42 and DGRP-508 showed decreases of approximately 0.8 mg and 0.3 mg, respectively (Fig. 3). Together, these results indicate that mass at eclosion responded to treatment, but the direction and magnitude of the response depended strongly on genetic background. Estimated marginal means values and standard deviations for body mass across treatments and genetic backgrounds are provided in Supplemental Table S5.

### 3.2. Reproduction

F_1_ fertility. A total of 29,668 flies were counted to measure F_1_ fertility across treatments and genetic backgrounds (Table 2). For consistency, offspring produced after day 7 were excluded from the analysis (Fig.4; Supplemental Table S1). F_1_ fertility was influenced by oxygen availability, temperature, genetic background, and the interaction between oxygen and temperature (Table 3). Across lines, hypoxia generally reduced F_1_ fertility relative to control, with mean declines of approximately 1.2 to 2.7 offspring per female over 7 days. Low temperature alone had a smaller and less consistent effect, with several lines showing fertility similar to, or slightly higher than, control. The combined treatment produced the strongest reduction in fertility across all genetic backgrounds, exceeding the effects of either stressor alone. The magnitude of this reduction varied among lines. DGRP-42 had the lowest mean F_1_ fertility under the combined treatment, averaging 4.45 offspring per female, about 50% lower than control. DGRP-391 and DGRP-491 also showed strong reductions under the combined treatment, of ∼52% and ∼32%, respectively. DGRP-57 had an intermediate reduction of about ∼37% relative to control. DGRP-508, retained the highest fertility under the combined treatment, averaging 6.10 offspring per female, although this still represented a decline relative to control. Overall, the combined treatment reduced F_1_ fertility by ∼35–55% across lines, corresponding to declines of approximately 3 to mora than 5 offspring per female.

**Figure 4:**
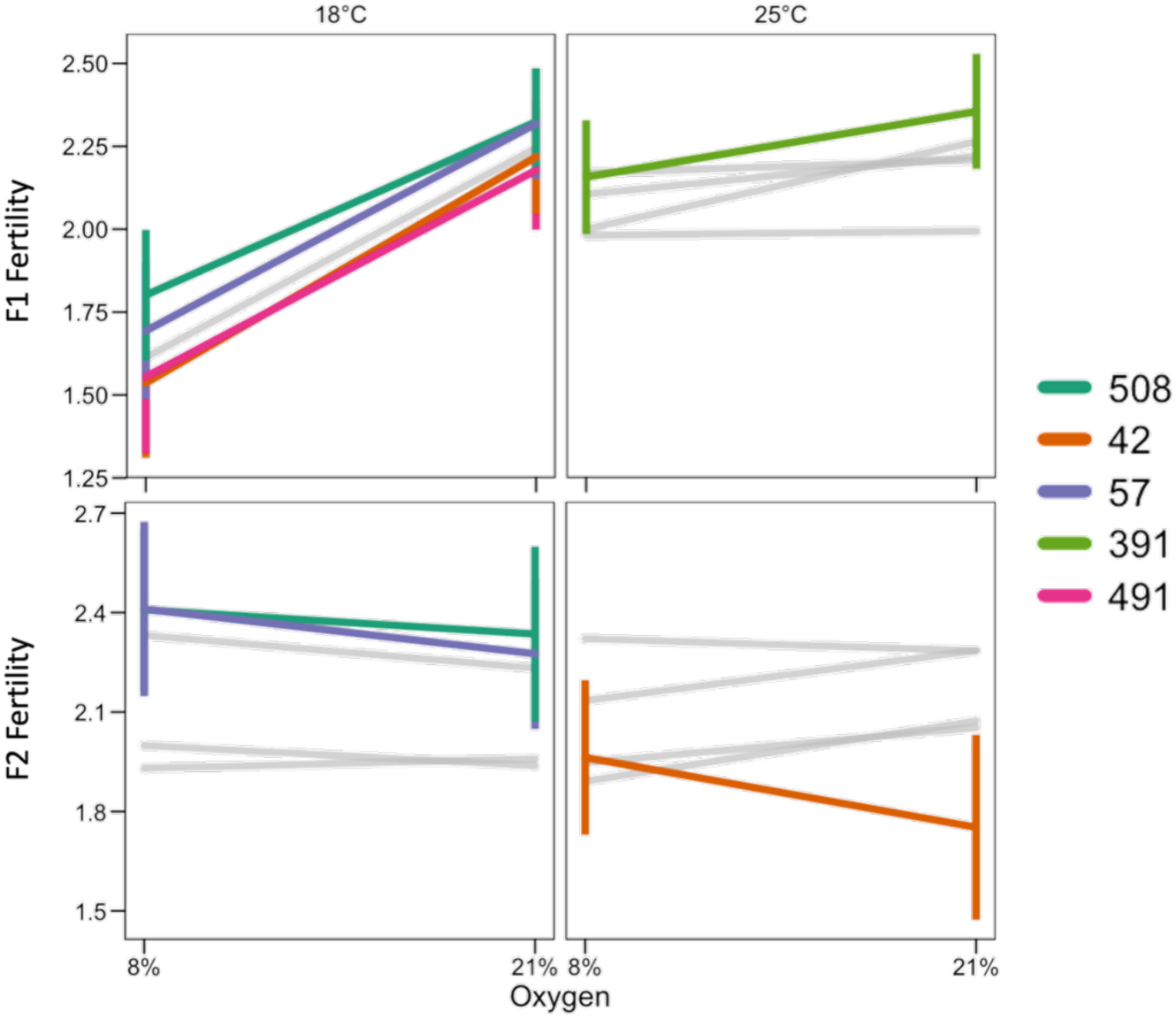
Reaction norm of the estimates of cross-generational fertility for both temperature treatments across two oxygen levels. Stocks are color-coded, with non-gray lines indicating statistical significance from the post-hoc analysis.

**Table 2:** Summary of F_1_ fertility per female over 7 days by genetic background, at the different treatments. Means and number of females used to calculate F_1_ fertility.

| *Mean / n* | *Control Temp* | *Low Temp* | *Hypoxia* | *Combined* |
| --- | --- | --- | --- | --- |
| 391 | Mean=9.82 | 9 | 7.15 | 4.72 |
|  | N=156 | 181 | 197 | 172 |
| 42 | 9 | 8.31 | 6.63 | 4.52 |
|  | 189 | 179 | 179 | 185 |
| 491 | 6.66 | 8.38 | 6.66 | 4.55 |
|  | 167 | 174 | 163 | 178 |
| 508 | 8.43 | 9.95 | 7.76 | 6.1 |
|  | 140 | 164 | 186 | 189 |
| 57 | 8.38 | 9.42 | 7.18 | 5.25 |
|  | 160 | 183 | 151 | 194 |

**Table 3:**
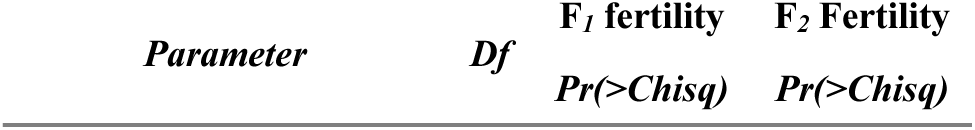
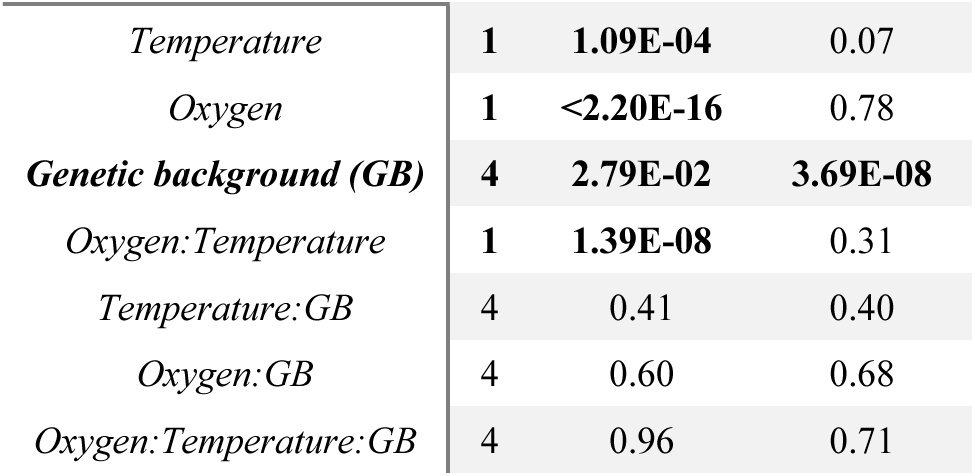
Mixed-effects model results for the effects of oxygen, temperature, genetic background, and their interactions on F_1_ and F_2_ fertility.

| *Parameter* | *Df* | *F<sub>1</sub> fertility* | *F<sub>2</sub> Fertility* |
| --- | --- | --- | --- |
|  |  | *Pr(>Chisq)* | *Pr(>Chisq)* |
| *Temperature* | **1** | **1.09E-04** | 0.07 |
| *Oxygen* | **1** | **<2.20E-16** | 0.78 |
| ***Genetic background (GB)*** | **4** | **2.79E-02** | **3.69E-08** |
| *Oxygen:Temperature* | **1** | **1.39E-08** | 0.31 |
| *Temperature:GB* | 4 | 0.41 | 0.40 |
| *Oxygen:GB* | 4 | 0.60 | 0.68 |
| *Oxygen:Temperature:GB* | 4 | 0.96 | 0.71 |

**Table 4.** Summary of F_2_ fertility per female over 7 days by genetic background, at the different treatments. Means and number of females used to calculate F_2_ fertility.

| **GB** | ***Control*** | ***Low Temp*** | ***Hypoxia*** | ***Combined*** |
| --- | --- | --- | --- | --- |
| 391 | 9.1 | 7.4 | 7.3 | 7.8 |
|  | 96 | 62 | 48 | 70 |
| 42 | 6.5 | 7.7 | 7.5 | 8.1 |
|  | 44 | 56 | 64 | 68 |
| 491 | 9.5 | 10 | 8 | 10.8 |
|  | 46 | 67 | 38 | 49 |
| 508 | 10.3 | 11.2 | 10.7 | 11.7 |
|  | 60 | 44 | 44 | 52 |
| 57 | 10.4 | 10.4 | 8.9 | 11.5 |
|  | 52 | 65 | 46 | 44 |

F_2_ Fertility. A total of 118,061 flies were scored to quantify F_2_ fertility. As for F_1_ fertility, counts beyond day 7 were excluded. Mean F_2_ fertility differed among genetic backgrounds, spanning approximately 6.5 to 11.7 offspring per female across treatments _2_ fertility (Table 3), but neither oxygen availability, temperature, nor their interaction produced a consistent overall effect.

Although treatment effects were not significant overall, some line-specific differences were evident. DGRP-391 showed its highest mean F_2_ fertility under control conditions, and its lowest under hypoxia, corresponding to a reduction of ∼1.8 offspring per female. In contrast, DGRP-42 showed its lowest F_2_ fertility in the control treatment and its highest in the combined treatment, an increase of ∼1.6 offspring per female. DGRP-491 and DGRP-57 also had higher F_2_ fertility in the combined treatment than under hypoxia, with increases of approximately 1.3 and 1.5 offspring per female, respectively. DGRP-508 had the highest F_2_ fertility across all treatments, peaking under the combined treatment at 11.7 offspring per female, about ∼1.4 higher than in control conditions. Overall, treatment-related changes in F_2_ fertility were modest, generally around 1 to 2 offspring per female, and varied in direction among genetic backgrounds.

Oogenesis. A total of 453 ovaries were dissected and stained for analyses of oocyte counts and cell death (Supplemental Table S1). DGRP-57 was excluded due to insufficient sample size. The intermediate time point, 2 days post-eclosion, was also excluded because ovaries at that age contained oocytes spanning all developmental stages without a clear dominant stage group, limiting stage-specific comparisons (n=126).

The number of oocytes was strongly influenced by developmental stage, which showed the largest effect in the model (p<2.2×10^−16^). Oocyte number also differed significantly by time point (p=0.0014), temperature (p=1.04×10^−7^), and genetic background (p<0.001; Table 5). Oxygen availability did not have a significant overall effect, but treatment responses depended on the interaction among oxygen availability, genetic background, temperature, and timepoint (Table 5). Consistent with the structured nature of oogenesis, differences due to treatment were higher when examined within specific stages and time points. In DGRP-491 at timepoint 0, hypoxia was associated with substantially higher predicted mean oocyte numbers than normoxia in early stages of oogenesis. In stages S1–7, the predicted mean increased from 1.28 under normoxia to 3.57 under hypoxia, and in the germarium it increased from 1.10 to 3.39. These differences were not maintained at timepoint 5, however, where stages S1–7 showed nearly identical predicted means under normoxia and hypoxia, 3.84 vs 3.82, respectively.

**Figure 5:**
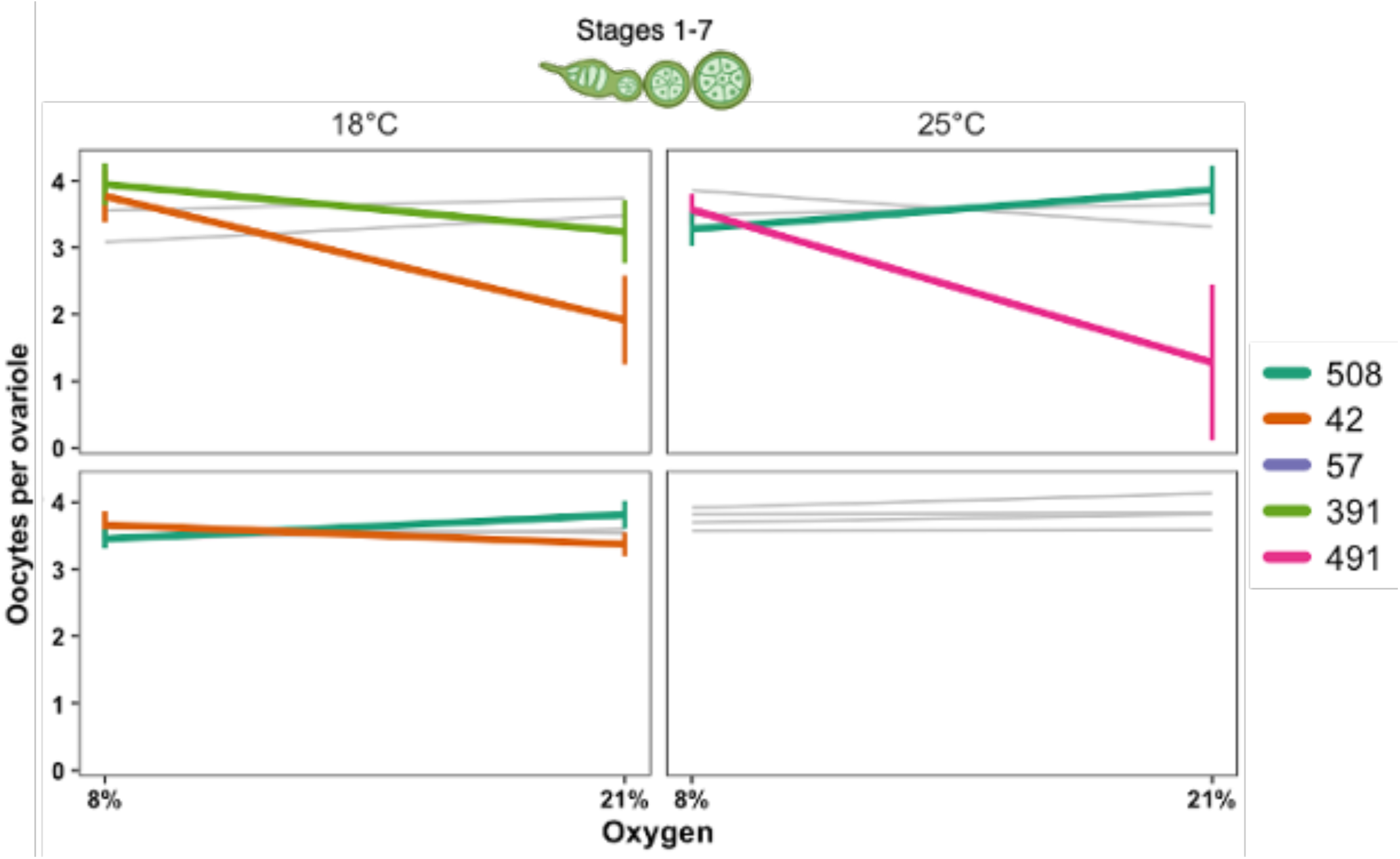
Mean number of oocytes for developmental stages S1-7 across two oxygen treatments, two temperatures, and two time points (0 and 5 days post-eclosion) for each genetic background. Significant genotypes for a given oxygen-temperature combination are highlighted in color, while non-significant responses are shown in grey. Asterisks denote significant contrast between oxygen treatments withing genotype.

**Table 5:** Mixed-effects model results for the effects of oxygen, temperature, genetic background, and their interactions on number of oocytes.

| *Parameter* | *Df* | *Pr(>Chisq)* |
| --- | --- | --- |
| *Stage* | 4 | **<2.20E-16** |
| *Timepoint* | 1 | **1.43E-03** |
| *Temperature* | 1 | **1.04E-07** |
| *Genetic background (GB)* | 3 | **3.98E-04** |
| *Oxygen* | 1 | 0.20 |
| *Temperature:GB* | 3 | **2.88E-02** |
| *GB:Oxygen* | 3 | **1.79E-03** |
| *Temperature:Oxygen* | 1 | 0.30 |
| *Timepoint:GB* | 3 | 0.73 |
| *Timepoint:Temperature* | 1 | 0.10 |
| *Timepoint:Oxygen* | 1 | **7.40E-02** |
| *Temperature:GB:Oxygen* | 3 | **6.12E-03** |
| *Timepoint:Temperature:Oxygen* | 1 | 0.21 |
| *Timepoint:Temperature:GB* | 3 | **1.67E-03** |
| *Timepoint:GB:Oxygen* | 3 | **1.77E-04** |
| *Timepoint:Temperature:GB:Oxygen* | 3 | **5.74E-05** |

A similarly pattern was observed in DGRP-42 at low temperature and time point 0. In this line, several stage groupings had higher predicted mean oocyte numbers under hypoxia than under normoxia. For instance, in stages S8–1 the predicted mean increased from 0.51 to 2.36, and in stages S12–14 it increased from 1.22 to 3.07. As in DGRP-491, these differences were substantially reduced by time point 5.

Not all lines responded in the same direction. In DGRP-508, predicted mean oocyte numbers in stages S1–7 tended to be lower under hypoxia than under normoxia at time point 0 under both temperature conditions.

Under control temperature, the predicted mean decreased from 3.86 to 3.28 under hypoxia, and under low temperature it decreased from 3.48 to 3.08 in the combined treatment. Together, these results show that treatment effects on oogenesis were strongly dependent on developmental stage, time point, and genetic background.

**Figure 6:**
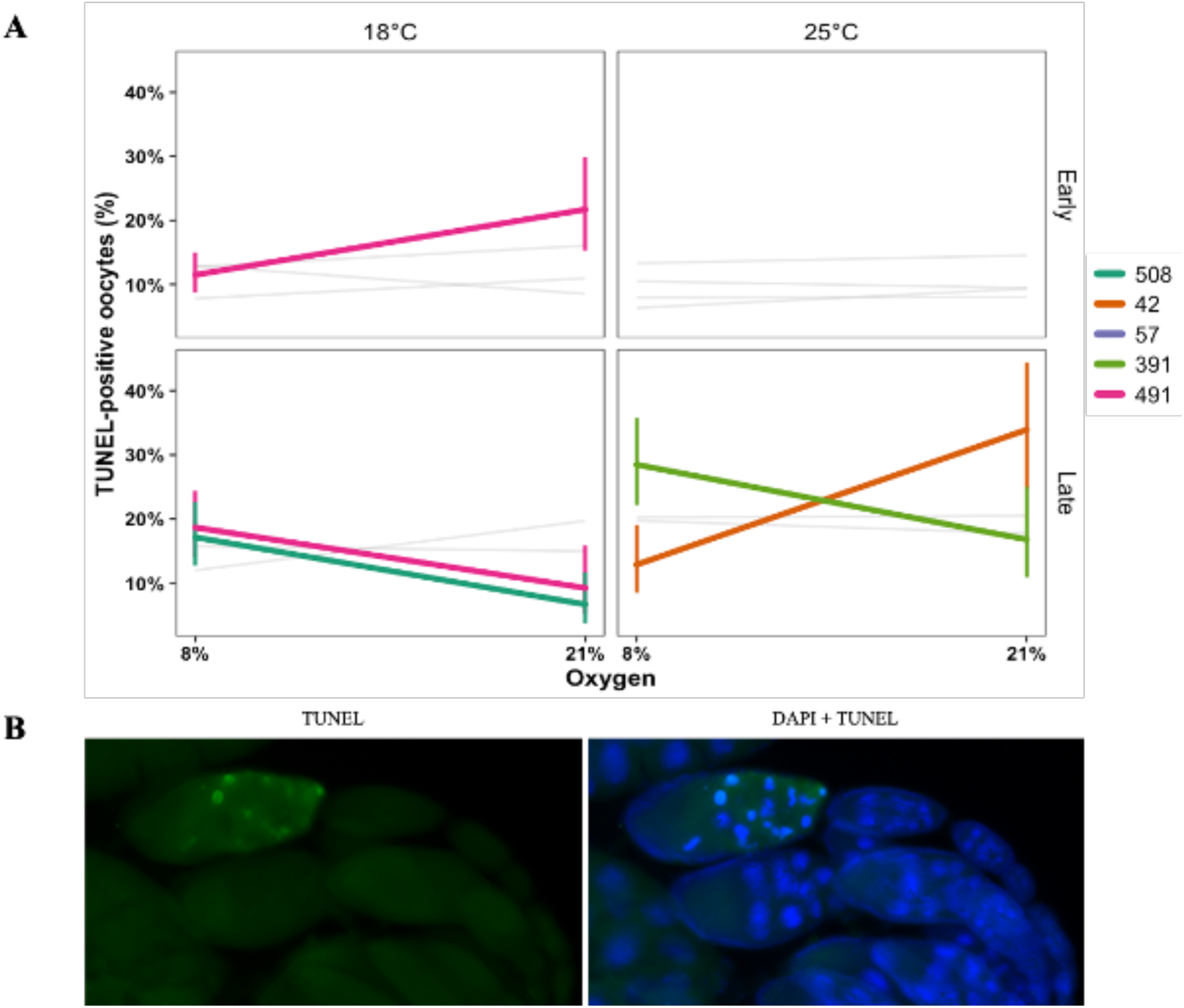
(A) Reaction norms of estimated TUNEL-positive oocytes per ovariole, averaged across timepoints, shown for early (germarium + S1–7) and late (S8–14) oogenic stages under two temperature treatments and two oxygen levels. Lines represent genetic backgrounds. Colored lines indicate genotypes with a significant oxygen contrast within a given temperature and stage, while non-significant responses are shown in grey. (B) Representative image of oocytes stained with TUNEL (green) and TUNEL + DAPI (blue), highlighting cell death.

Cell death. The proportion of TUNEL-positive oocytes differed strongly between early and late oogenesis (p<2.2×10^−16^; Table 6), with substantially higher and more variable cell death in late stages. Time point and genetic background also affected cell death. In contrast, temperature and oxygen availability did not have significant overall main effects, but their effects depended on developmental stage and genotype. Significant interactions were detected between stage and temperature (p<1×10^−9^), stage and oxygen (p=0.003), and the four-way interactions among stage, oxygen, temperature, and genetic background (Table 6).

**Table 6:** Type II Wald χ² tests for fixed effects from the binomial generalized linear mixed-effects model of oocyte cell death. The response was modeled as the number of TUNEL-positive and TUNEL-negative oocytes. Note here that the germarium and S1-7 are grouped into a single early stage and the other three stages are grouped as a single late stage.

| *Parameter* | *Df* | *Pr(>Chisq)* |
| --- | --- | --- |
| *Stage* | 1 | **<2.20E-16** |
| *Timepoint* | 1 | **1.96E-04** |
| *Temperature* | 1 | 0.42 |
| *Genetic background (GB)* | 3 | **8.63E-03** |
| *Oxygen* | 1 | 0.175 |
| *Temperature:GB* | 3 | 0.652 |
| *GB:Oxygen* | 3 | 0.308 |
| *Temperature:Oxygen* | 1 | 0.759 |
| *Stage:GB* | 3 | **5.13E-03** |
| *Stage:Temperature* | 1 | **6.44E-10** |
| *Stage:Oxygen* | 1 | **3.08E-03** |
| *Temperature:GB:Oxygen* | 3 | 0.311 |
| *Stage:Temperature:Oxygen* | 1 | **2.82E-02** |
| *Stage:Temperature:GB* | 3 | 0.85 |
| *Stage:GB:Oxygen* | 3 | **1.46E-10** |
| *Stage:Temperature:GB:Oxygen* | 3 | **1.50E-03** |

During early oogenesis, the proportion of TUNEL-positive oocytes was generally low across genetic backgrounds. DGRP-491 showed one of the clearest early stage responses: relative to the control treatment, cell death was reduced under low temperature by 1.28 but increased under hypoxia by 2.14. DGRP-508 also showed low early-stage cell death across treatments, with the strongest reduction occurring under the combined treatment, 2.69 lower than control, representing the lowest predicted cell-death levels observed during early oogenesis among all genotypes.

Late oogenesis showed substantially higher and more variable levels of cell death. DGRP-508 exhibited one of the strongest late-stage responses: under low temperature, cell death increased by 2.64, whereas under the combined treatment it was reduced by 1.40, creating one of the largest contrasts observed across all developmental stages. DGRP-491 also showed marked late-stage responses, with higher cell death under the combined treatment, 2.29, than the control treatment, 1.36. DGRP-391 showed variation as well, but without the extreme shifts observed in DGRP-508 or DGRP-491. Overall, these results indicate that treatment effects on oocyte cell death were concentrated in late oogenesis and varied strongly among genetic backgrounds.

## 4. Discussion

Understanding how organisms respond to environmental stressors is critical for predicting their tolerance and adaptability in changing ecosystems. While many studies investigate the effects of individual stressors, natural environments often present organisms with multiple simultaneous challenges that interact in complex ways. These interactions can lead to synergistic, antagonistic, or additive effects, which may significantly alter physiological and reproductive outcomes. In this study, we explored the combined effects of hypoxia and low temperature on physiological traits, fertility, and reproduction in *Drosophila melanogaster* across five different genetic backgrounds. By examining responses to both single and combined stressors, we aimed to better understand how genetic variation shapes reproductive and physiological responses and identify the unique challenges posed by multiple environmental pressures.

### 4.1 Physiological and reproductive stress response is influenced by genetic background

The results confirm a genetic influence on the variation in metabolic rate, thermal tolerance, and body mass under hypoxia and low temperature. Across most genetic backgrounds, hypoxia generally led to increased respiratory quotient (RQ) values, indicating a shift in metabolic pathways likely toward carbohydrate metabolism. This aligns with findings from Van Voorhies (2009), who reported increasing RQ values in *Drosophila* under hypoxic conditions. However, this study also noted exceptions in certain genotypes, such as the ones observed in this study for DGRP-57 and DGRP-491, which exhibited minimal changes. This suggests that genetic diversity among individuals may lead to differential responses, a phenomenon also observed in other ectothermic species (Hoffmann et al., 2013) where some genotypes also demonstrated metabolic tolerance under stress.

The thermal tolerance, measured by CT_max_, was significantly influenced by genetic background, with interactions between temperature and oxygen driving some of the observed changes in this trait. Most genotypes exhibited increased CT_max_ under hypoxia, particularly at control temperature, suggesting a potential stress-buffering effect of low oxygen on heat tolerance. This observation follows the patterns described previously for *Drosophila melanogaster* (C Teague et al., 2017), where a positive correlation between hypoxia and thermal tolerance was described. Which could indicate that in certain genetic backgrounds, hypoxia may confer an advantage in extreme heat scenarios, highlighting the need for further research into species-specific responses to simultaneous thermal and oxygen stress.

Body mass at eclosion was strongly influenced by genetic background and environmental conditions, with hypoxia often associated with increased mass relative to normoxia. This pattern contrasts with the well-established negative relationship between oxygen availability and structural body size reported in Drosophila and other insects (Czarnoleski et al., 2023; Heinrich et al., 2011; Noguchi et al., 2022). Importantly, most previous studies have quantified body size using morphological traits such as wing area or pupal case dimensions, whereas the present study measured whole-body mass immediately following eclosion.

Body mass at this developmental stage may reflect a short-term physiological state rather than fixed morphological size, as it can be influenced by hydration status, gut contents, and allocation of internal reserves. Newly eclosed flies also undergo rapid physiological adjustments, including meconium excretion, which can introduce additional variability in mass measurements. Consequently, the observed increases in mass under hypoxia should not be interpreted as evidence that low oxygen promotes increased structural body size.

These results suggest that hypoxia and temperature may alter patterns of mass allocation or physiological condition depending on the genetic background (C. Teague et al., 2017). Differences among genetic backgrounds indicate that some genotypes may respond to hypoxic development by retaining greater internal reserves or water content, whereas others show reduced mass under the same conditions. Future studies combining morphological size measurements with mass and compositional analyses would help to better understand these genetic differences. In particular, measuring dry mass alongside wet mass would help disentangle structural size changes from differences in water content or internal reserves, providing a clearer physiological interpretation of the body mass responses observed here.

### 4.2 F_1_ fertility is Dramatically Reduced Under Combined Stress

Hypoxia negatively impacted F_1_ fertility across all genetic backgrounds, consistent with existing evidence that low oxygen disrupts metabolic and physiological functions critical for survival (Callier et al., 2015; Lee et al., 2019; Lighton, 2007; Peck & Maddrell, 2005; Polan et al., 2020; Rand et al., 2018; Romero et al., 2007; Van Voorhies, 2009; Zhou et al., 2007). In contrast, low temperature alone did not exhibit a significant effect, suggesting that *Drosophila* may tolerate cold better than hypoxia in isolation. However, when combined with hypoxia, low temperature further reduced F_1_ fertility, highlighting the compounding effects of multiple stressors. The interaction between oxygen and temperature was significant, revealing a possible combined effect of these environmental factors.

Comparing F_1_ and F_2_ fertility across genotypes, we observed notable differences in reproductive outcome. DGRP-491 exhibited consistently lower F_1_ fertility across treatments but showed a relative recovery in F_2_ fertility, suggesting that while the first generation suffered immediate reproductive costs under stress, the second generation was less affected. In contrast, DGRP-391, which maintained higher F_1_ fertility, exhibited a slight decline in F_2_, indicating potential delayed effects of stress exposure. DGRP-42, DGRP-57, and DGRP-508, which performed poorly in F_1_ fertility, continued to show low reproductive success in F_2_, suggesting an inability to recover from early environmental challenges. These results suggest genotype-specific differences in reproductive recovery, where some lines show improved output in the second generation while others remain sensitive. However, because all F2 crosses were reared at 25°C under normoxia, these patterns could reflect either true maternal or epigenetic carryover effects or residual differences in parental condition; the present design does not allow these mechanisms to be distinguished. While we did not observe a direct effect of low temperature on F_1_ fertility, previous studies have reported temperature effects on reproductive success across generations. For example, Sgrò & Hoffmann (1998) measure fecundity, whereas our study quantified the number of adult offspring produced during the first 8 days post-mating. Differences between studies may therefore reflect both assays duration and the reproductive metric measured.

### 4.3 Differential Oogenesis Response by Time Point and Treatment

The effects of hypoxia and low temperature on oogenesis were highly variable across genetic backgrounds, time point, and developmental stages. At the early time point (0 days post-eclosion), hypoxia had an effect on the oocyte number across genetic backgrounds, indicating that initial reproductive investment might be sensitive to oxygen availability to some genetic backgrounds. Similar genetic background variation in reproductive responses to hypoxia has been documented in Drosophila, where oxygen limitation can differentially affect ovarian activity and egg production depending on genetic background and physiology (Arquier et al., 2006; Wingrove & O’Farrell, 1999). For example, the combined treatment appeared to stimulate oogenesis across most genetic backgrounds except DGRP-491, which again exhibited a contrasting response. These results suggest that response mechanisms may vary when more than one stressor is applied and is influenced by genetic background, supporting the need to consider genetic variation and the effect of interactions between more than one stressor when assessing reproductive sensitivity under stress.

Later in development (5 days post-eclosion), oocyte number often shifted in both magnitude and direction relative to early measurements, indicating that duration of exposure and reproductive age modulate oogenesis responses to environmental stress. In several genetic backgrounds, hypoxia was associated with increased oocyte numbers, consistent with compensatory or delayed oogenesis following prolonged exposure to stress (Canal Domenech & Fricke, 2023; Meena et al., 2024; Rivera-Rincón et al., 2024). However, when hypoxia was combined with low temperature, oocyte production was frequently reduced, highlighting that combined stressors impose stronger constraints on reproductive output than either stressor alone. Importantly, these patterns were not uniform across genotypes, with some lines exhibiting increased oocyte numbers under combined stress while others showed pronounced reductions. Together, these results demonstrate that total oocyte production reflects a dynamic balance between environmental conditions, reproductive timing, and genetic background, rather than a fixed response to hypoxia or temperature alone.

### 4.4 Cell Death as an Indicator of Reproductive Cost

The TUNEL assay data provided further insight into the influence of genetic background on reproductive costs associated with stress. Early oogenesis (germarium and stages 1–7) was characterized by generally low levels of cell death across treatments, indicating substantial buffering of early oocyte development. This relative protection of early oogenic stages is consistent with previous work showing that early oocyte development is often maintained under stress to preserve future reproductive potential (Arquier et al., 2006). Nevertheless, a subset of genetic backgrounds exhibited pronounced oxygen and temperature effects on cell death at early stages, suggesting that early oocyte survival can be selectively sensitive to combined stressors in certain genotypes.

In contrast, late oogenesis (stages 8–14) showed a major vulnerability to environmental changes. Several genetic backgrounds, most notably DGRP-508 and DGRP-491, exhibited large changes in late-stage oocyte cell death due to temperature, oxygen availability and their interaction. Increased cell death during late oogenesis under hypoxic or combined stress conditions is consistent with the idea that apoptosis acts as a regulatory mechanism to limit reproductive investment when energetic or physiological constraints are exceeded (Drummond-Barbosa & Spradling, 2001; Pritchett et al., 2009). In other backgrounds, such as DGRP-42, elevated late-stage apoptosis under hypoxia and combined treatments, suggests higher sensitivity to oxygen limitation during the final oocyte maturation. These findings indicate that reproductive costs of environmental stress are disproportionately expressed during late oogenesis, when energetic investment is highest and opportunities for compensation are reduced.

### 4.5 Genotype specific responses to individual and combined stress

Our results offer new perspectives into how genetic diversity shapes the ability to tolerate multiple environmental stressors, and we can directly relate these findings to prior studies on chill coma recovery and oxidative stress resistance (Mackay et al., 2012; Weber et al., 2012). By comparing the performance of each genotype across these assays, we can better understand their relative resilience and vulnerabilities to compounded stress.

DGRP-42, DGRP-57, and DGRP-508 consistently performed poorly across nearly all assays, both individually and under combined stress. These lines had the lowest survival rates in the paraquat-induced oxidative stress assay and the slowest chill coma recovery times in prior studies (Mackay et al., 2012; Weber et al., 2012), and this trend was reflected in our study consistent with a conserved genetic architecture for these two stressors (Zhao and Haddad 2012). Physiologically, DGRP-42 exhibited a significant increase in respiratory quotient (RQ) under hypoxia, indicating a shift toward carbohydrate metabolism, which was further exacerbated by low temperature. In terms of reproductive output, DGRP-42 showed the most pronounced reduction in fertility under the combined stressors, consistent with its poor performance across stress assays. The oogenesis data further reinforced these findings, with DGRP-42 showing a significant reduction in oocyte counts at 5 days post-eclosion under combined stress, as well as increased TUNEL-positive oocytes, suggesting higher levels of apoptosis in reproductive tissues. These findings suggest that DGRP-42 is particularly sensitive to hypoxia and low-temperature stress, with limited capacity for compensatory mechanisms, such as increased oogenesis or reduced apoptosis, under these conditions.

DGRP-57, despite demonstrating the lowest survival in the paraquat assay and poor chill coma recovery, showed a more complex response. Under hypoxia, DGRP-57 exhibited a lower body mass at eclosion compared to many other lines. However, the reproductive output was still severely impacted by hypoxia, with reduced fertility across both stress treatments. The oogenesis assays showed minimal impacts at the earlier time points, but DGRP-57 did not exhibit the same dramatic decrease in oocyte counts at 5 days post-eclosion as DGRP-42. However, it did show higher TUNEL-positive oocytes under hypoxia, DGRP-508, followed in our metrics the patterns previously observed, which showed low survival in oxidative stress and slow recovery from chill coma. Physiologically, DGRP-508 exhibited a significant increase in RQ under hypoxia, with a reduced body mass at eclosion compared to controls, consistent with a potentially lower metabolic adaptation to stress. In terms of reproductive outcome, DGRP-508 showed consistently low fertility across all stress treatments, and its oogenesis response was not significantly different from DGRP-42. This line also demonstrated higher TUNEL-positive oocytes, indicating increased apoptosis. The overall performance of DGRP-508 suggests that it is highly sensitive to combined hypoxia and low-temperature stress, with limited capacity to respond through metabolic or reproductive changes.

In contrast, DGRP-391 and DGRP-491 showed greater tolerance across most assays. DGRP-391, which had higher survival rates in the paraquat assay and a faster recovery from chill coma (Mackay et al., 2012; Weber et al., 2012), demonstrated relatively high reproductive outcome under hypoxia and cold stress, as well as physiological responses that were more tolerant to the effects of combined stressors. Under hypoxia, DGRP-391 showed a minimal shift in RQ, suggesting that it was able to maintain metabolic stability. While its fertility did decrease under combined stress, it remained higher than any of the other genotypes, and oogenesis assays revealed relatively stable oocyte counts at both early and later time points. TUNEL-positive oocytes in DGRP-391 were also lower compared to the other lines, indicating a reduced level of apoptosis in the ovaries. These results suggest that DGRP-391 is well-adapted to buffer the combined effects of hypoxia and cold stress, with a strong capacity for both physiological and reproductive tolerance.

DGRP-491 showed similarly strong tolerance across stressors. This line, which also exhibited higher survival rates in paraquat and faster chill coma recovery (Mackay et al., 2012; Weber et al., 2012), demonstrated a remarkable ability to recover in terms of reproductive output in the second generation (F_2_). While DGRP-491 exhibited lower fertility in the first generation (F_1_) under combined stress, its F_2_ fertility was significantly higher, suggesting possible transgenerational resilience or maternal effects, though the present design cannot distinguish between these mechanisms because all F2 crosses were reared under standard conditions. Metabolically, DGRP-491 showed minimal changes in RQ under hypoxia and a stable body mass at eclosion, indicating strong physiological regulation. In the oogenesis assays, DGRP-491 maintained a relatively high number of oocytes at 5 days post-eclosion, and the TUNEL-positive oocytes was lower compared to other lines, reinforcing its tolerance under combined stress. These findings suggest that DGRP-491 can not only tolerate immediate stress effects but also recover reproductively in subsequent generations.

## 5. Conclusions

These findings underscore the significant role of genetic background in shaping stress responses, particularly when multiple stressors are involved. The observed changes in metabolic pathways, such as higher respiratory quotient (RQ) values under hypoxia, suggest that certain genotypes may adaptively adjust to low oxygen levels, while others, like DGRP-491, show limited metabolic changes, indicating varying levels of tolerance to environmental stress. Similarly, thermal tolerance was found to be influenced by genetic background and the interplay between oxygen levels and temperature. Some genotypes exhibited increased tolerance to heat under hypoxia, suggesting that low oxygen may help buffer stress in extreme heat conditions.

Fertility under combined hypoxia and low temperature in the F_1_ generation revealed that while low temperature alone was not significantly impacted, the combination of both stressors magnified the effects, particularly in genotypes like DGRP-491, which showed consistent susceptibility to stress. Fertility in the F_2_ generation further emphasized the importance of genetic background, with some genotypes maintaining high reproductive output under stress, while others experienced declines, particularly when exposed to both hypoxia and low temperature. Oogenesis progression showed varied responses across genotypes, with hypoxia affecting oocyte counts in a genotype- and time point manner. Early developmental stages saw reduced oocyte counts for most genotypes, but certain lines, such as DGRP-508, demonstrated adaptive responses. This variation highlights the need to consider genetic diversity when assessing reproductive tolerance under environmental stress. Additionally, the TUNEL assay results provided insights into the reproductive costs of stress, showing that hypoxia influenced apoptosis in a genotype-specific manner, prioritizing survival over reproduction in stress-sensitive lines. Overall, this study enhances our understanding of how environmental changes involving multiple stressors interact with genetic factors to affect physiological and reproductive traits. These findings have broader implications for understanding the adaptive capacity of natural populations and underscore the importance of further research into the genetic and molecular mechanisms driving these responses, including the roles of stress-response proteins and gene expression changes.

## Acknowledgments

We would like to thank, Dr. Graff, Dr. Koehler, and Dr. Watanabe at the College of Veterinary Medicine, at Auburn University for allowing us to use the BZ-X fluorescence microscope and their mentorship in microscopy. We would like to thank Dr. Hood, Dr. Schwartz, Dr. Bernal, and Dr. Stoeckel for their revisions and feedback on the manuscript. We would like to thank members of the Stevison Lab for helpful feedback throughout this project. Research reported in this publication was supported by the National Institute of General Medical Sciences of the National Institutes of Health under Award Number R35GM147501. The content is solely the responsibility of the authors and does not necessarily represent the official views of the National Institutes of Health.

## Bibliography

Arking, R., Buck, S., Wells, R. A., & Pretzlaff, R. (1988). Metabolic rates in genetically based long lived strains of Drosophila. Exp Gerontol, 23(1), 59–76. doi:10.1016/0531-5565(88)90020-4

Arquier, N., Vigne, P., Duplan, E., Hsu, T., Therond, P., Frelin, C., & D’Angelo, G. (2006). Analysis of the hypoxia-sensing pathway in Drosophila melanogaster - pubmed. The Biochemical journal, 393(Pt 2). doi:10.1042/BJ20050675

Bates, D., Mächler, M., Bolker, B., & Walker, S. (2014). Fitting linear mixed-effects models using lme4. arXiv preprint arXiv:1406.5823.

Boyd, P. W., Collins, S., Dupont, S., Fabricius, K., Gattuso, J.-P., Havenhand, J., … Pörtner, H.-O. (2018). Experimental strategies to assess the biological ramifications of multiple drivers of global ocean change—a review. Global Change Biology, 24(6). doi:10.1111/gcb.14102

Callier, V., Hand, S. C., Campbell, J. B., Biddulph, T., & Harrison, J. F. (2015). Developmental changes in hypoxic exposure and responses to anoxia in Drosophila melanogaster. J Exp Biol, 218(Pt 18), 2927–2934. doi:10.1242/jeb.125849

Canal Domenech, B., & Fricke, C. (2023). Developmental heat stress interrupts spermatogenesis inducing early male sterility in Drosophila melanogaster - pubmed. Journal of Thermal Biology, 114. doi:10.1016/j.jtherbio.2023.103589

Chiu, C.-L., Clack, N., & community, t. n. (2022). Napari: A python multi-dimensional image viewer platform for the research community | microscopy and microanalysis | cambridge core. Microscopy and Microanalysis, 28(S1). doi:10.1017/S1431927622006328

Crain, C. M., Kroeker, K., & Halpern, B. S. (2008). Interactive and cumulative effects of multiple human stressors in marine systems. Ecology Letters, 11(12). doi:10.1111/j.1461-0248.2008.01253.x

Czarnoleski, M., Szlachcic, E., Privalova, V., Labecka, A. M., Sikorska, A., Sobczyk, Ł., … Angilletta Jr, M. J. (2023). Oxygen and temperature affect cell sizes differently among tissues and between sexes of Drosophila melanogaster. Journal of insect physiology, 150, 104559.

Dang, D., Le, M., Irmer, T., Angay, O., Fichtl, B., & Schwarz, B. (2021). Apeer: An interactive cloud platform for microscopists to easily deploy deep learning. Zenodo.

DeVries, Z. C., & Appel, A. G. (2013). Standard metabolic rates of lepisma saccharina and thermobia domestica: Effects of temperature and mass. J Insect Physiol, 59(6), 638–645. doi:10.1016/j.jinsphys.2013.04.002

Drummond-Barbosa, D., & Spradling, A. C. (2001). Stem cells and their progeny respond to nutritional changes during Drosophila oogenesis. Developmental biology, 231(1), 265–278.

Flatt, T. (2011). Survival costs of reproduction in Drosophila. Exp Gerontol, 46(5), 369–375. doi:10.1016/j.exger.2010.10.008

Folguera, G., Bastías, D. A., Caers, J., Rojas, J. M., Piulachs, M.-D., Bellés, X., & Bozinovic, F. (2011). An experimental test of the role of environmental temperature variability on ectotherm molecular, physiological and life-history traits: Implications for global warming. Comparative Biochemistry and Physiology Part A: Molecular & Integrative Physiology, 159(3), 242–246.

Fox, J., Weisberg, S., Adler, D., Bates, D., Baud-Bovy, G., Ellison, S., … Graves, S. (2012). Package ‘car’. Vienna: R Foundation for Statistical Computing, 16.

Gorr, T., Gassmann, M., & Wappner, P. (2006). Sensing and responding to hypoxia via hif in model invertebrates. Journal of insect physiology, 52(4). doi:10.1016/j.jinsphys.2006.01.002

Gorr, T. A., Tomita, T., Wappner, P., & Bunn, H. F. (2004). Regulation of Drosophila hypoxia-inducible factor (hif) activity in sl2 cells. Journal of Biological Chemistry, 279(34), 36048–36058. doi:10.1074/jbc.M405077200

Gunderson, A. R., Dillon, M. E., & Stillman, J. H. (2017). Estimating the benefits of plasticity in ectotherm heat tolerance under natural thermal variability. Functional Ecology, 31(8), 1529–1539.

Gunderson, A. R., & Stillman, J. H. (2015). Plasticity in thermal tolerance has limited potential to buffer ectotherms from global warming. Proceedings of the Royal Society B: Biological Sciences, 282(1808), 20150401.

Heinrich, E., Farzin, M., Klok, C., & Harrison, J. (2011). The effect of developmental stage on the sensitivity of cell and body size to hypoxia in Drosophila melanogaster. The Journal of Experimental Biology, 214(9). doi:10.1242/jeb.051904

Helmuth, B., Russell, B. D., Connell, S. D., Dong, Y., Harley, C. D., Lima, F. P., … Mieszkowska, N. (2014). Beyond long-term averages: Making biological sense of a rapidly changing world. Climate Change Responses 2014 1:1, 1(1). doi:10.1186/s40665-014-0006-0

Hoffmann, A. A., Chown, S. L., & Clusella-Trullas, S. (2013). Upper thermal limits in terrestrial ectotherms: How constrained are they? Functional Ecology, 27(4), 934–949.

Hoffmann, A. A., Dagher, H., Hercus, M. J., & Berrigan, D. (1997). Comparing different measures of heat resistance in selected lines of *Drosophila melanogaster*. Journal of insect physiology, 43(4), 393–405.

Hoffmann, A. A., & Sgrò, C. M. (2011). Climate change and evolutionary adaptation. Nature, 470(7335), 479–485.

Hu, X. P., & Appel, A. G. (2004). Seasonal variation of critical thermal limits and temperature tolerance in formosan and eastern subterranean termites (isoptera: Rhinotermitidae). Environmental entomology, 33(2), 197–205.

Huang, W., Massouras, A., Inoue, Y., Peiffer, J., Ràmia, M., Tarone, A. M., … Mackay, T. F. (2014). Natural variation in genome architecture among 205 Drosophila melanogaster genetic reference panel lines. Genome Res, 24(7), 1193–1208. doi:10.1101/gr.171546.113

Hulbert, A. J., Clancy, D. J., Mair, W., Braeckman, B. P., Gems, D., & Partridge, L. (2004). Metabolic rate is not reduced by dietary-restriction or by lowered insulin/igf-1 signalling and is not correlated with individual lifespan in Drosophila melanogaster. Experimental Gerontology, 39(8), 1137–1143.

Jia, D., Xu, Q., Xie, Q., Mio, W., & Deng, W. M. (2016). Automatic stage identification of Drosophila egg chamber based on dapi images. Sci Rep, 6(1), 18850. doi:10.1038/srep18850

Kassambara, A., & Kassambara, M. A. (2020). Package ‘ggpubr’. R package version 0.1, 6(0).

Kluyver, T., Ragan-Kelley, B., Perez, F., Granger, B., Bussonnier, M., … Team, J. D. (2016). Jupyter notebooks – a publishing format for reproducible computational workflows. doi:10.3233/978-1-61499-649-1-87

Lazzaro, B. P., Flores, H. A., Lorigan, J. G., & Yourth, C. P. (2008). Genotype-by-environment interactions and adaptation to local temperature affect immunity and fecundity in *Drosophila melanogaster*. PLOS Pathogens, 4(3). doi:10.1371/journal.ppat.1000025

Lee, B., Barretto, E. C., Grewal, S. S., Lee, B., Barretto, E. C., & Grewal, S. S. (2019). Torc1 modulation in adipose tissue is required for organismal adaptation to hypoxia in Drosophila. Nature Communications 2019 10:1, 10(1). doi:10.1038/s41467-019-09643-7

Lenth, R., & Lenth, M. R. (2018). Package ‘lsmeans’. The American Statistician, 34(4), 216–221.

Lighton, J. R. B. (2007). Hot hypoxic flies: Whole-organism interactions between hypoxic and thermal stressors in Drosophila melanogaster. Journal of Thermal Biology, 32(3), 134–143. doi:10.1016/j.jtherbio.2007.01.009

Lord, J. S., Leyland, R., Haines, L. R., Barreaux, A. M. G., Bonsall, M. B., Torr, S. J., & English, S. (2021). Effects of maternal age and stress on offspring quality in a viviparous fly. Ecology Letters, 24(10). doi:10.1111/ele.13839

Mackay, T. F., Richards, S., Stone, E. A., Barbadilla, A., Ayroles, J. F., Zhu, D., … Gibbs, R. A. (2012). The Drosophila melanogaster genetic reference panel. Nature, 482(7384), 173–178. doi:10.1038/nature10811

Meehan, T. L., Yalonetskaya, A., Joudi, T. F., & McCall, K. (2015). Detection of cell death and phagocytosis in the Drosophila ovary. Methods Mol Biol, 1328, 191–206. doi:10.1007/978-1-4939-2851-4_14

Meena, A., Nardo, A. N. D., Maggu, K., Sbilordo, S. H., Roy, J., Snook, R. R., & Lüpold, S. (2024). Fertility loss and recovery dynamics after repeated heat stress across life stages in male Drosophila melanogaster: Patterns and processes. Royal Society Open Science, 11(10). doi:10.1098/rsos.241082

Meiselman, M. R., Kingan, T. G., & Adams, M. E. (2018). Stress-induced reproductive arrest in Drosophila occurs through eth deficiency-mediated suppression of oogenesis and ovulation. BMC Biology, 16(1). doi:10.1186/s12915-018-0484-9

Mockett, R. J., Orr, W. C., Rahmandar, J. J., Sohal, B. H., & Sohal, R. S. (2001). Antioxidant status and stress resistance in long- and short-lived lines of Drosophila melanogaster. Exp Gerontol, 36(3), 441–463. doi:10.1016/s0531-5565(00)00258-8

Noguchi, K., Yokozeki, K., Tanaka, Y., Suzuki, Y., Nakajima, K., Nishimura, T., & Goda, N. (2022). Sima, a Drosophila homolog of hif-1α, in fat body tissue inhibits larval body growth by inducing tribbles gene expression. Genes to Cells, 27(2). doi:10.1111/gtc.12913

Peck, L. S., & Maddrell, S. H. P. (2005). Limitation of size by hypoxia in the fruit fly *Drosophila melanogaster*. Journal of Experimental Zoology Part A: Comparative Experimental Biology, 303A(11). doi:10.1002/jez.a.211

Piggott, J., Lange, K., Townsend, C., & Matthaei, C. (2012). Multiple stressors in agricultural streams: A mesocosm study of interactions among raised water temperature, sediment addition and nutrient enrichment - pubmed. PLoS One, 7(11). doi:10.1371/journal.pone.0049873

Polan, D. M., Alansari, M., Lee, B., & Grewal, S. S. (2020). Early-life hypoxia alters adult physiology and reduces stress resistance and lifespan in Drosophila. J Exp Biol, 223(Pt 22), jeb.226027. doi:10.1242/jeb.226027

Pritchett, T. L., Tanner, E. A., & McCall, K. (2009). Cracking open cell death in the Drosophila ovary. Apoptosis : an international journal on programmed cell death, 14(8), 969–979. doi:10.1007/s10495-009-0369-z

Promislow, D. E., & Haselkorn, T. S. (2002). Age-specific metabolic rates and mortality rates in the genus Drosophila. Aging cell, 1(1), 66–74. doi:10.1046/j.1474-9728.2002.00009.x

Rand, D. M., Mossman, J. A., Zhu, L., Biancani, L. M., & Ge, J. Y. (2018). Mitonuclear epistasis, genotype-by-environment interactions, and personalized genomics of complex traits in Drosophila. IUBMB Life, 70(12). doi:10.1002/iub.1954

Rivera-Rincón, N., Altindag, U. H., Amin, R., Graze, R. M., Appel, A. G., & Stevison, L. S. (2024). “A comparison of thermal stress response between Drosophila melanogaster and Drosophila pseudoobscura reveals differences between species and sexes”. Journal of insect physiology, 153. doi:10.1016/j.jinsphys.2024.104616

Romero, N. M., Dekanty, A., & Wappner, P. (2007). Cellular and developmental adaptations to hypoxia: A Drosophila perspective. In Methods in enzymology (Vol. 435, pp. 123–144): Elsevier.

Schindelin, J., Arganda-Carreras, I., Frise, E., Kaynig, V., Longair, M., Pietzsch, T., … Cardona, A. (2012). Fiji: An open-source platform for biological-image analysis. Nat Methods, 9(7), 676–682. doi:10.1038/nmeth.2019

Sgrò, C. M., & Hoffmann, A. A. (1998). Heritable variation for fecundity in field-collected *Drosophila melanogaster* and their offspring reared under different environmental temperatures. Evolution, 52(1). doi:10.1111/j.1558-5646.1998.tb05146.x

Shiran, M. G., Bailey, N. P., McCann, L., Rivera-Rincón, N., Saurette, E., & Stevison, L. S. (2026). Efficient rflp-based protocol for routine authentication of Drosophila. microPublication Biology, 2026, 10.17912/micropub.biology.001949.

Singh, N. D. (2019). Wolbachia infection associated with increased recombination in Drosophila. G3 (Bethesda), 9(1), 229–237. doi:10.1534/g3.118.200827

Sponsler, R., & Appel, A. (1991). Temperature tolerances of the formosan and eastern subterranean termites (isoptera: Rhinotermitidae). Journal of Thermal Biology, 16(1), 41–44.

Teague, C., Youngblood, J., Ragan, K., AngillettaJr, M., & VandenBrooks, J. (2017). A positive genetic correlation between hypoxia tolerance and heat tolerance supports a controversial theory of heat stress. Biol Lett, 13(11). doi:10.1098/rsbl.2017.0309

Teague, C., Youngblood, J. P., Ragan, K., Angilletta, M. J., Jr., & VandenBrooks, J. M. (2017). A positive genetic correlation between hypoxia tolerance and heat tolerance supports a controversial theory of heat stress. Biol Lett, 13(11). doi:10.1098/rsbl.2017.0309

Teder, T., & Kaasik, A. (2023). Early-life food stress hits females harder than males in insects: A meta-analysis of sex differences in environmental sensitivity. Ecology Letters, 26(8). doi:10.1111/ele.14241

Valko, A., Perez-Pandolfo, S., Sorianello, E., Brech, A., Wappner, P., & Melani, M. (2022). Adaptation to hypoxia in Drosophila melanogaster requires autophagy - pubmed. Autophagy, 18(4). doi:10.1080/15548627.2021.1991191

Van der Walt, S., Schönberger, J. L., Nunez-Iglesias, J., Boulogne, F., Warner, J. D., Yager, N., … Yu, T. (2014). Scikit-image: Image processing in python. PeerJ, 2, e453.

Van Voorhies, W. A. (2009). Metabolic function in Drosophila melanogaster in response to hypoxia and pure oxygen. J Exp Biol, 212(19), 3132–3141. doi:10.1242/jeb.031179

Van Voorhies, W. A., Khazaeli, A. A., & Curtsinger, J. W. (2003). Selected contribution: Long-lived Drosophila melanogaster lines exhibit normal metabolic rates. J Appl Physiol (1985), 95(6), 2605–2613; discussion 2604. doi:10.1152/japplphysiol.00448.2003

Van Voorhies, W. A., Khazaeli, A. A., & Curtsinger, J. W. (2004). Testing the “rate of living” model: Further evidence that longevity and metabolic rate are not inversely correlated in Drosophila melanogaster. Journal of Applied Physiology, 97(5), 1915–1922.

Weber, A. L., Khan, G. F., Magwire, M. M., Tabor, C. L., Mackay, T. F., & Anholt, R. R. (2012). Genome-wide association analysis of oxidative stress resistance in Drosophila melanogaster. PLoS One, 7(4), e34745. doi:10.1371/journal.pone.0034745

Wickham, H. (2011). Ggplot2. Wiley interdisciplinary reviews: computational statistics, 3(2), 180–185.

Wingrove, J., & O’Farrell, P. (1999). Nitric oxide contributes to behavioral, cellular, and developmental responses to low oxygen in Drosophila - pubmed. Cell, 98(1). doi:10.1016/S0092-8674(00)80610-8

Zhou, D., Xue, J., Chen, J., Morcillo, P., Lambert, J. D., White, K. P., & Haddad, G. G. (2007). Experimental selection for Drosophila survival in extremely low o(2) environment. PLoS One, 2(5), e490. doi:10.1371/journal.pone.0000490

Zizzari, Z. V., & Ellers, J. (2014). Rapid shift in thermal resistance between generations through maternal heat exposure. Oikos, 123(11), 1365–1370.

