## Supplement for "Effects of hypoxia and low temperature on female physiology and reproduction of *Drosophila melanogaster*"

**Supplemental material**

**Figure S1:** Density plots of the distribution of the selected lines for paraquat on the left and chill coma on the right. Specifically, we selected the lines 42, 57, 391, 491, and 508 which had paraquat survival values of 20.69, 12.34, 18.05, 25.86 and 12.57 and chill coma recovery times of 29.07, 24.59, 12.74, 11.93, and 29.21 minutes, respectively. These lines were obtained from the Bloomington Drosophila Stock Center (NIH P40OD018537).

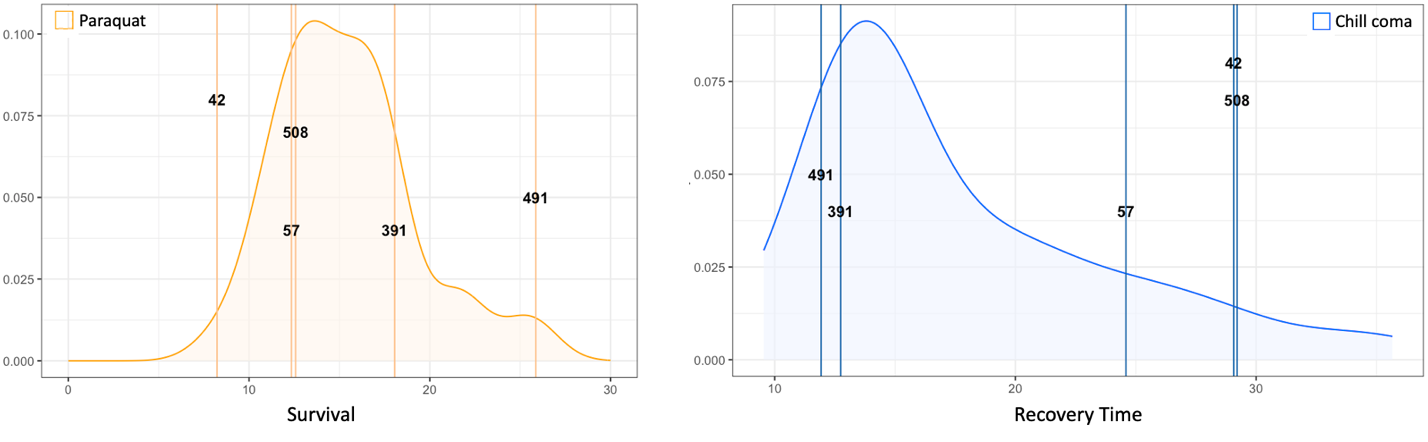

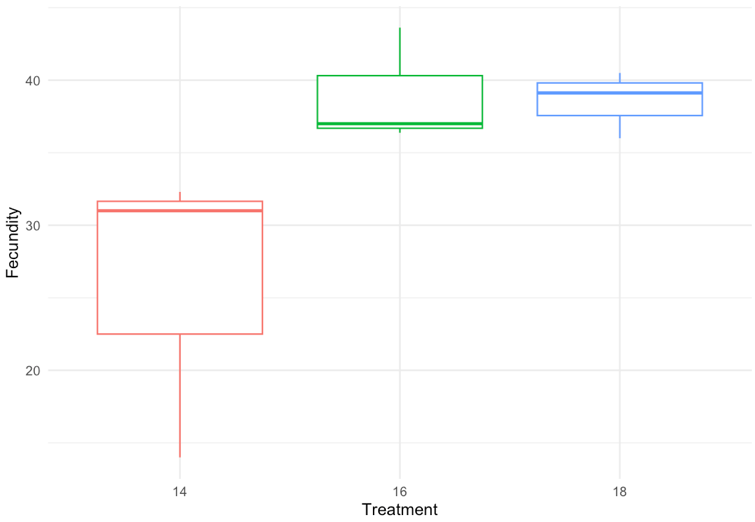

**Figure S2:** Fecundity per female over 8 days for DGRP-217 at three different temperatures, 14 ℃, 16 ℃, and 18 ℃.

**Table S1:** Sample sizes for each assay across treatments and genetic backgrounds (GBs)

| **Treatment** | **Genetic Background** | **Survivorship** | **RQ** | **CT_max_** | **Body mass** | **Fecundity** | **Oogenesis** |
| --- | --- | --- | --- | --- | --- | --- | --- |
| Control | *42* | *189* | *8* | *7* | *10* | *44* | *23* |
| ColdTemp | *42* | *179* | *13* | *16* | *10* | *56* | *18* |
| Hypoxia | *42* | *179* | *6* | *6* | *10* | *64* | *12* |
| Combined | *42* | *185* | *11* | *17* | *9* | *68* | *18* |
| Control | *57* | *160* | *12* | *10* | *9* | *52* | *17* |
| ColdTemp | *57* | *183* | *12* | *10* | *10* | *65* | *24* |
| Hypoxia | *57* | *151* | *8* | *7* | *9* | *46* | *23* |
| Combined | *57* | *194* | *10* | *11* | *9* | *44* | *32* |
| Control | *391* | *156* | *0* | *0* | *10* | *96* | *28* |
| ColdTemp | *391* | *181* | *15* | *16* | *10* | *62* | *20* |
| Hypoxia | *391* | *197* | *5* | *4* | *10* | *48* | *18* |
| Combined | *391* | *172* | *19* | *15* | *10* | *70* | *25* |
| Control | *491* | *167* | *6* | *7* | *10* | *46* | *21* |
| ColdTemp | *491* | *174* | *6* | *6* | *10* | *67* | *22* |
| Hypoxia | *491* | *163* | *6* | *7* | *11* | *38* | *19* |
| Combined | *491* | *178* | *15* | *16* | *10* | *49* | *39* |
| Control | *508* | *140* | *3* | *3* | *10* | *60* | *11* |
| ColdTemp | *508* | *164* | *6* | *7* | *10* | *44* | *19* |
| Hypoxia | *508* | *186* | *10* | *10* | *10* | *44* | *19* |
| Combined | *508* | *189* | *12* | *11* | *9* | *52* | *45* |

**Table S2:** Estimated marginal means for VCO_2_ showing the effects of **stock, temperature, and oxygen treatment** on the response variable with their associated standard errors and confidence intervals where applicable.

| **Genetic Background** | **Temperature** | **Oxygen** | **emmean** | **SE** | **df** | **lower.CL** | **upper.CL** |
| --- | --- | --- | --- | --- | --- | --- | --- |
| 42 | Control | Control | 0.539 | 0.0497 | 22.3 | 0.436 | 0.642 |
| 491 | Control | Control | 0.372 | 0.0555 | 28.3 | 0.258 | 0.485 |
| 508 | Control | Control | 0.446 | 0.073 | 26.4 | 0.296 | 0.596 |
| 57 | Control | Control | 0.515 | 0.0377 | 26.9 | 0.438 | 0.592 |
| 42 | Low | Control | 0.562 | 0.0396 | 28.7 | 0.481 | 0.643 |
| 491 | Low | Control | 0.461 | 0.0429 | 46.6 | 0.374 | 0.547 |
| 508 | Low | Control | 0.438 | 0.0382 | 30.9 | 0.361 | 0.516 |
| 57 | Low | Control | 0.493 | 0.0412 | 22.9 | 0.407 | 0.578 |
| 42 | Control | Low | 0.512 | 0.0558 | 36.2 | 0.398 | 0.625 |
| 491 | Control | Low | 0.4 | 0.0468 | 36.5 | 0.305 | 0.494 |
| 508 | Control | Low | 0.481 | 0.0516 | 26.4 | 0.375 | 0.587 |
| 57 | Control | Low | 0.467 | 0.0524 | 27 | 0.359 | 0.574 |
| 42 | Low | Low | 0.456 | 0.036 | 34.3 | 0.383 | 0.529 |
| 491 | Low | Low | 0.403 | 0.0412 | 22.1 | 0.318 | 0.489 |
| 508 | Low | Low | 0.462 | 0.0397 | 29.5 | 0.381 | 0.543 |
| 57 | Low | Low | 0.453 | 0.0389 | 30.9 | 0.374 | 0.532 |

**Table S3:** Estimated marginal means for VO_2_ showing the effects of **stock, temperature, and oxygen treatment** on the response variable with their associated standard errors and confidence intervals where applicable.

| **Genetic Background** | **Temperature** | **Oxygen** | **emmean** | **SE** | **df** | **lower.CL** | **upper.CL** |
| --- | --- | --- | --- | --- | --- | --- | --- |
| 42 | Control | Control | 0.614 | 0.0778 | 24.7 | 0.453 | 0.774 |
| 491 | Control | Control | 0.446 | 0.0848 | 31.1 | 0.273 | 0.619 |
| 508 | Control | Control | 0.589 | 0.113 | 27.6 | 0.357 | 0.82 |
| 57 | Control | Control | 0.601 | 0.058 | 28.9 | 0.482 | 0.719 |
| 42 | Low | Control | 0.66 | 0.0603 | 31.7 | 0.537 | 0.783 |
| 491 | Low | Control | 0.535 | 0.0636 | 42.5 | 0.407 | 0.664 |
| 508 | Low | Control | 0.514 | 0.0583 | 31.1 | 0.395 | 0.633 |
| 57 | Low | Control | 0.539 | 0.0643 | 25.5 | 0.406 | 0.671 |
| 42 | Control | Low | 0.563 | 0.0843 | 34.4 | 0.391 | 0.734 |
| 491 | Control | Low | 0.498 | 0.0704 | 35.7 | 0.355 | 0.64 |
| 508 | Control | Low | 0.62 | 0.0798 | 27.6 | 0.456 | 0.783 |
| 57 | Control | Low | 0.466 | 0.0807 | 28.4 | 0.301 | 0.631 |
| 42 | Low | Low | 0.56 | 0.0544 | 34.6 | 0.45 | 0.671 |
| 491 | Low | Low | 0.456 | 0.0643 | 25.2 | 0.323 | 0.588 |
| 508 | Low | Low | 0.555 | 0.0604 | 32.1 | 0.432 | 0.678 |
| 57 | Low | Low | 0.495 | 0.0593 | 31.8 | 0.374 | 0.616 |

**Table S4:** Estimated marginal means for *CT_max_* showing the effects of **stock, temperature, and oxygen treatment** on the response variable with their associated standard errors and confidence intervals where applicable.

| **Genetic Background** | **Temperature** | **Oxygen** | **emmean** | **SE** | **df** | **lower.CL** | **upper.CL** |
| --- | --- | --- | --- | --- | --- | --- | --- |
| 42 | Control | Control | 38.9 | 0.215 | 20.9 | 38.4 | 39.3 |
| 42 | Control | Low | 39 | 0.224 | 25 | 38.5 | 39.4 |
| 42 | Low | Control | 39.2 | 0.159 | 19.8 | 38.8 | 39.5 |
| 42 | Low | Low | 38.4 | 0.142 | 20.9 | 38.1 | 38.7 |
| 491 | Control | Control | 39.6 | 0.235 | 17.8 | 39.2 | 40.1 |
| 491 | Control | Low | 38.9 | 0.203 | 31.1 | 38.5 | 39.3 |
| 491 | Low | Control | 39.1 | 0.202 | 55.8 | 38.7 | 39.5 |
| 491 | Low | Low | 39.2 | 0.198 | 12.5 | 38.8 | 39.6 |
| 508 | Control | Control | 38.7 | 0.317 | 25 | 38 | 39.3 |
| 508 | Control | Low | 39.1 | 0.215 | 20.9 | 38.6 | 39.5 |
| 508 | Low | Control | 39.3 | 0.173 | 28.7 | 39 | 39.7 |
| 508 | Low | Low | 39.3 | 0.195 | 19.8 | 38.9 | 39.7 |
| 57 | Control | Control | 38.9 | 0.177 | 33.2 | 38.5 | 39.3 |
| 57 | Control | Low | 39.2 | 0.215 | 20.9 | 38.8 | 39.7 |
| 57 | Low | Control | 39.3 | 0.195 | 19.8 | 38.9 | 39.7 |
| 57 | Low | Low | 39.1 | 0.162 | 32.5 | 38.7 | 39.4 |

**Table S5:** Estimated marginal means for body mass showing the effects of **stock, temperature, and oxygen treatment** on the response variable with their associated standard errors and confidence intervals where applicable.

| **Genetic Background** | **Temperature** | **Oxygen** | **emmean** | **SE** | **df** | **lower.CL** | **upper.CL** |
| --- | --- | --- | --- | --- | --- | --- | --- |
| 42 | Control | Control | 0.00664 | 8.35E-05 | 140 | 0.00647 | 0.00681 |
| 42 | Control | Low | 0.00575 | 8.35E-05 | 140 | 0.00558 | 0.00592 |
| 42 | Low | Control | 0.00641 | 8.35E-05 | 140 | 0.00624 | 0.00658 |
| 42 | Low | Low | 0.00683 | 8.80E-05 | 140 | 0.00666 | 0.00701 |
| 491 | Control | Control | 0.00575 | 8.35E-05 | 140 | 0.00558 | 0.00592 |
| 491 | Control | Low | 0.00624 | 7.96E-05 | 140 | 0.00608 | 0.00639 |
| 491 | Low | Control | 0.00516 | 8.35E-05 | 140 | 0.00499 | 0.00533 |
| 491 | Low | Low | 0.00676 | 8.35E-05 | 140 | 0.00659 | 0.00693 |
| 508 | Control | Control | 0.00727 | 8.35E-05 | 140 | 0.0071 | 0.00744 |
| 508 | Control | Low | 0.00697 | 8.35E-05 | 140 | 0.0068 | 0.00714 |
| 508 | Low | Control | 0.00601 | 8.35E-05 | 140 | 0.00584 | 0.00618 |
| 508 | Low | Low | 0.00694 | 8.80E-05 | 140 | 0.00677 | 0.00712 |
| 57 | Control | Control | 0.0063 | 8.80E-05 | 140 | 0.00613 | 0.00647 |
| 57 | Control | Low | 0.0067 | 8.80E-05 | 140 | 0.00653 | 0.00687 |
| 57 | Low | Control | 0.00562 | 8.35E-05 | 140 | 0.00545 | 0.00579 |
| 57 | Low | Low | 0.00624 | 8.80E-05 | 140 | 0.00607 | 0.00642 |
